# Fractional Deletion Of Compound Kushen Injection, A Natural Compound Mixture, Indicates Cytokine Signaling Pathways Are Critical For Its Perturbation Of The Cell Cycle

**DOI:** 10.1101/462135

**Authors:** TN Aung, S Nourmohammadi, Z Qu, Y Harata-Lee, J Cui, HY Shen, AJ Yool, T Pukala, Du Hong, RD Kortschak, DL Adelson

## Abstract

We have used computational and experimental biology approaches to identify candidate mechanisms of action of a traditional Chinese medicine. Compound Kushen Injection (CKI), in a breast cancer cell line in which CKI causes apoptosis. Because CKI is a complex mixture of plant secondary metabolites, we used a high-performance liquid chromatography (HPLC) fractionation and reconstitution approach to define chemical fractions required for CKI to induce apoptosis in MDA-MB-231 cells. Our initial fractionation separated major from minor compounds, and showed that the major compounds accounted for little of the activity of CKI. By systematically perturbing the major compounds in CKI we found that removal of no single major compound could alter the effect of CKI on cell viability and apoptosis. However, simultaneous removal of two major compounds identified oxymatrine and oxysophocarpine as critical compounds with respect to CKI activity. We then used RNA sequencing and transcriptome analysis to correlate compound removal with gene expression and phenotype data. We determined that many compounds in CKI are required for its effectiveness in triggering apoptosis but that significant modulation of its activity is conferred by a small number of compounds. In conclusion, CKI may be typical of many plant based extracts that contain many compounds in that no single compound is responsible for all of the bioactivity of the mixture and that many compounds interact in a complex fashion to influence a network containing many targets.

## Introduction

Natural compounds are chemically diverse and have long served as resources for the identification of drugs (Harvey et al., 2015). However, the standard approach of fractionating natural product extracts to identify a single compound’s biological activity can fail because the original activity of the mixture is not present in single compounds after fractionation. This failure to identify single compounds implies that some natural product mixtures derive their activity from the interaction of several bioactive compounds within the mixture. Characterising the mode of action of natural product mixtures has remained a difficult task as the combinatorial complexity of such mixtures makes it unfeasible to screen all combinations of the compounds in the mixture.

We introduce here a “subtractive fractionation approach” using high performance liquid chromatography (HPLC) that can pinpoint significant interacting compounds within a mixture when coupled with a suitable bioassay. We combined this approach with RNAseq (RNA sequencing) characterisation of our bioassay, correlating the removal of interacting compounds with concomitant alterations in gene expression. This allows us to identify specific combinations of compounds associated with specific pathways and regulatory interactions. In this report, we have applied this approach for the first time to a particular Traditional Chinese Medicine (TCM) formulation: Compound Kushen Injection (CKI), which is used to treat approximately 30,000 cancer patients/day in China in conjunction with Western chemotherapy. CKI is composed primarily of alkaloids and flavonoids extracted from two herbal medicinal plants: Kushen (*Radix Sophorae Flavescentis*) and Baituling (*Rhizoma Smilacis Glabrae*). Twenty-one chromatographic peaks have been identified from CKI with eight compounds being recognized as major components on the basis of their abundance (Ma et al., 2014).

The extract containing the most abundant compounds in CKI is derived from Kushen herb which has a long history in the treatment of patients suffering immune function disorders (Xu et al., 2005;Cheng et al., 2006). The main component of CKI, macrozamin, is a derivative of baituling which has been a suggested therapeutic agent for the treatment of inflammatory disease (Jiang et al., 1997). Gao and colleagues showed that treatment with each of four of the main compounds of CKI (oxymatrine, matrine, sophoridine and N-methycytisine) at 4 mg/ml significantly decreased cell viability (Gao et al., 2018). However, these concentrations are relatively high when compared to the contributing concentration of these four main compounds in CKI (Ma et al., 2014). The two main components of CKI, matrine and oxymatrine, may have significant anticancer activities in various types of solid tumors including breast cancer non small lung cancer, cervical cancer, prostate cancer, synovial sarcoma, and hepatocellular carcinoma (Yu et al., 2009;Li et al., 2015;Wu et al., 2015;Cai et al., 2016;Wu et al., 2016;Aung et al., 2017;Gao et al., 2018;Zhou et al., 2018). In contrast, toxicity of medicinal herbs containing matrine and oxymatrine as main components has been reported (Wang and Yang, 2003). Administration of matrine 150 mg/kg and oxymatrine 360 mg/kg significantly increased cytochrome P450 family protein CYPB1/2 in rats demonstrating a potential therapeutic drawback of these two compounds (Yuan et al., 2010). Overall, understanding the effects of CKI based on the effects of single compounds present in CKI has been at best, partially successful.

Alternatively, by removing one, two or three compounds, we have been able to map the effects of these compounds and their interactions to effects on specific pathways based on altered gene expression profiles in a cell-based assay. This has illuminated the roles of several major compounds of CKI, which on their own have little or no activity in our bioassay. This approach can be used to dissect the roles and interactions of individual compounds from complex natural compound mixtures whose biological activity cannot be attributed to single purified compounds.

## Results

### Subtractive fractionation overview

Well resolved chromatographic separation of CKI was used to collect all of the major components of CKI as individual fractions (Figure 1A). We then reconstituted all of the separated fractions except for those we wished to subtract. We tested the reconstituted combination of compounds/peaks to see if removal of a single (N-1) or multiple compounds, (N-2 or N-3, where N represents the number of compounds in CKI), or removal of all major peaks (minor, MN) or depletion of all minor peaks (major, MJ) significantly altered the effect of CKI in our cell based assays. Our cell based assays (Qu et al., 2016) measured MDA-MB-231 (human breast adenocarcinoma) cell viability, cell cycle phase and cell apoptosis. A summary of the subtractive fractions used in the cell based assays is shown in Table 1. We then carried out RNA isolation of cells treated with CKI, individual compounds or CKI deletions for RNAseq. Differentially expressed (DE) genes in these samples allowed the association of specific compounds with cell phenotype and underlying alterations in gene regulation. By comparing DE genes across treatment combinations we identified specific candidate pathways that were altered by removal of single or multiple compounds, as detailed below.

**Figure 1:**
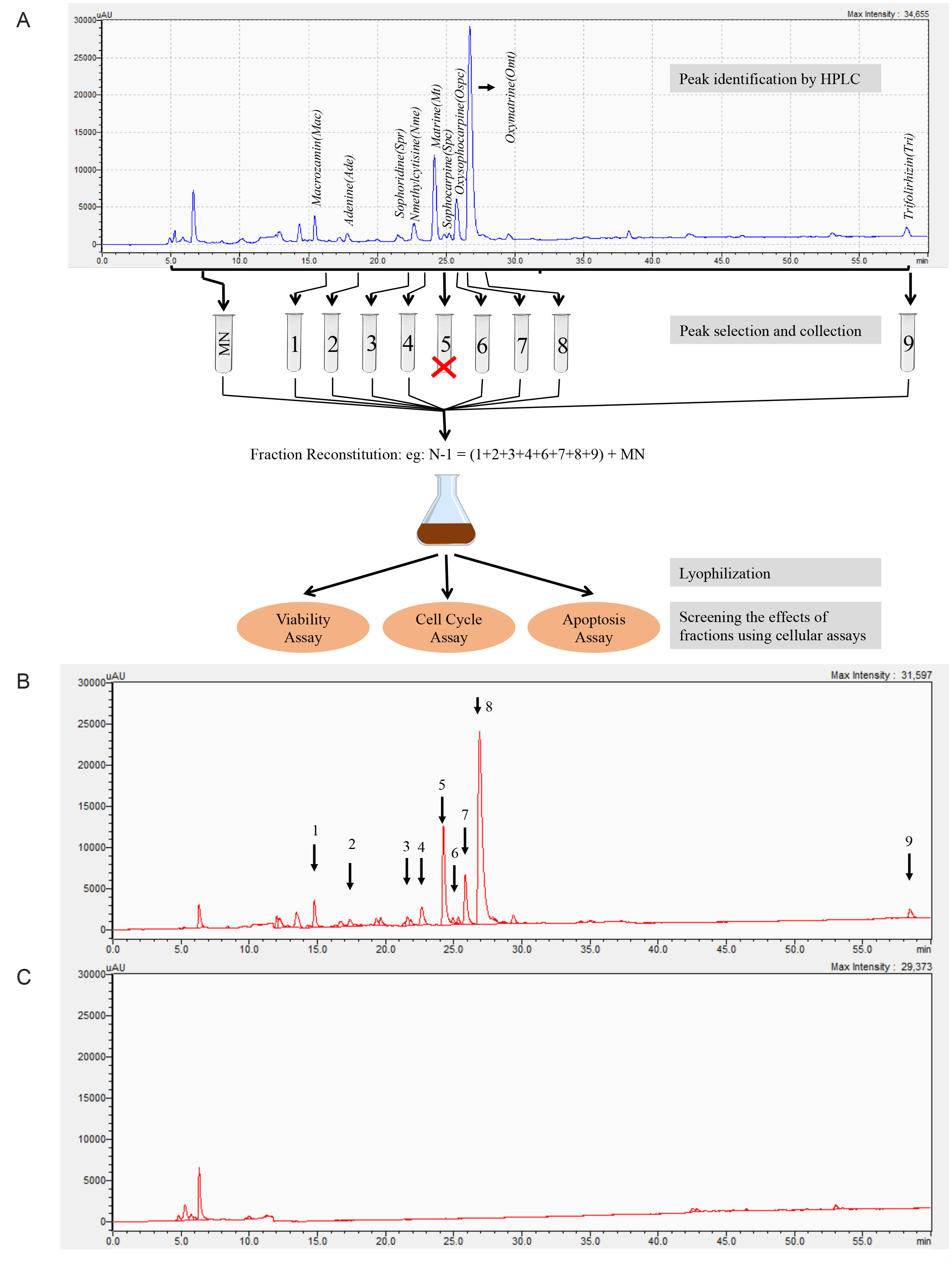
Fractionation of Compound Kushen Injection. (A) Diagram illustrating the process of subtractive fractionation, reconstitution, and screening of fractionated compounds using three cell-based assays. (B) HPLC profile of the 9 purified and reconstituted major peaks (MJ) demonstrating nine major compounds. (C) HPLC profile of reconstituted fractions not containing the 9 major compounds (MN) showing the remaining peaks with no remaining major compounds.

**Table 1:** Summarized results of HPLC fractionation and treatments using three cell-based assays. N represents the number of all compounds contained in CKI, Mac = macrozamin, Ade = adenine, Tri = trifolirhizin, Nme = N-methylcytisine, Spr = Sophoridine, Mt = Matrine, Omt = Oxymatrine, Spc = Sophocarpine, and Ospc = Oxysophocarpine. Significant results of CKI treatment were calculated based on UT whereas those of other treatments were calculated based on CKI treatments. Statistically significant results were represented as P<0.05 (*) or P<0.01 (**) P<0.001 (***), or P<0.0001 (****).

| HPLC Fractionation | Treatments | Proliferation Assay (MDA-MB-213) | Cell Cycle Assay (MDA-MB-213) | Apoptosis Assay in three cell lines |  |  |
| --- | --- | --- | --- | --- | --- | --- |
|  |  |  |  | MDA-MB-231 | HEK-293 | HFF |
| 9 known + small unknown (N) | Original CKI | * | * |  |  |  |
| 9known major compounds | MJ |  |  |  |  |  |
| Small unknown minor compounds | MN | * |  |  |  |  |
| N-1 | N-Mac |  |  |  |  |  |
|  | N-Ade |  |  |  |  |  |
|  | N-Tri |  |  |  |  |  |
|  | N-Nme |  |  |  |  |  |
|  | N-Spr |  |  |  |  |  |
|  | N-Mt |  |  |  |  |  |
|  | N-Omt |  |  |  |  |  |
|  | N-Ospc |  |  |  |  |  |
|  | N-Spc |  |  |  |  |  |
| N-3 | N-MacAdeTri |  |  |  |  |  |
|  | N-MtNmeSpr |  |  |  |  |  |
|  | N-OmtOspcSpc |  |  |  |  |  |
|  | N-MacOmtOspc | * | **** | **** | **** | ** |
| N-2 | N-MacAde |  |  |  |  |  |
|  | N-MacTri |  |  |  |  |  |
|  | N-AdeTri |  |  |  |  |  |
|  | N-MtNme |  |  |  |  |  |
|  | N-MtSpr |  |  |  |  |  |
|  | N-NmeSpr |  |  |  |  |  |
|  | N-OmtOspc | * | **** | **** | **** | ** |
|  | N-OmtSpc |  |  |  |  |  |
|  | N-OspcSpc |  |  |  |  |  |
| 3 compounds | MacAdeTri |  |  |  |  |  |
|  | MtNmeSpr |  |  |  |  |  |
|  | OmtOspcSpc |  |  |  |  |  |
|  | MacOmtOspc |  |  |  |  |  |
| 2 compounds | MacAde |  |  |  |  |  |
|  | MacTri |  |  |  |  |  |
|  | AdeTri |  |  |  |  |  |
|  | MtNme |  |  |  |  |  |
|  | MtSpr |  |  |  |  |  |
|  | NmeSpr |  |  |  |  |  |
|  | OmtOspc |  |  |  |  |  |
|  | OmtSpc |  |  |  |  |  |
|  | OspcSpc |  |  |  |  |  |

### HPLC fractions and content identification using LC-MS/MS

HPLC fractionation and reconstitution was used to generate a number of N-1, N-2, N-3, MJ and MN mixtures, (Figure 1A, B, C and Supplementary Figure 1) with specific combinations and their components shown in Table 1. The concentrations of known compounds in CKI and reconstituted subtractive fractions were determined from standard curves (Supplementary Data1) for nine available reference compounds, using cytisine as an internal standard (Table 2). The combined concentration of 9 reference compounds from CKI was approximately 10. 461 mg/ml, whereas subtractive fractions N-OmtOspc and N-MacOmtOspc had concentrations of reference compounds of 3.045 mg/ml and 2.335 mg/ml which were equivalent to the concentrations of these compounds in unfractionated CKI. The depleted OmtOspc and MacOmtOspc were not observed in the N-OmtOspc and N-MacOmtOspc respectively. These collectively suggested any effects observed after the treatments of N-OmtOspc and N-MacOmtOspc were not influenced by the concentrations. A total of 9 (N-1), 4 (N-3) and 9 (N-2) combinations, along with MJ and MN deletions were tested in our cell based assays (Table 1).

**Table 2:**
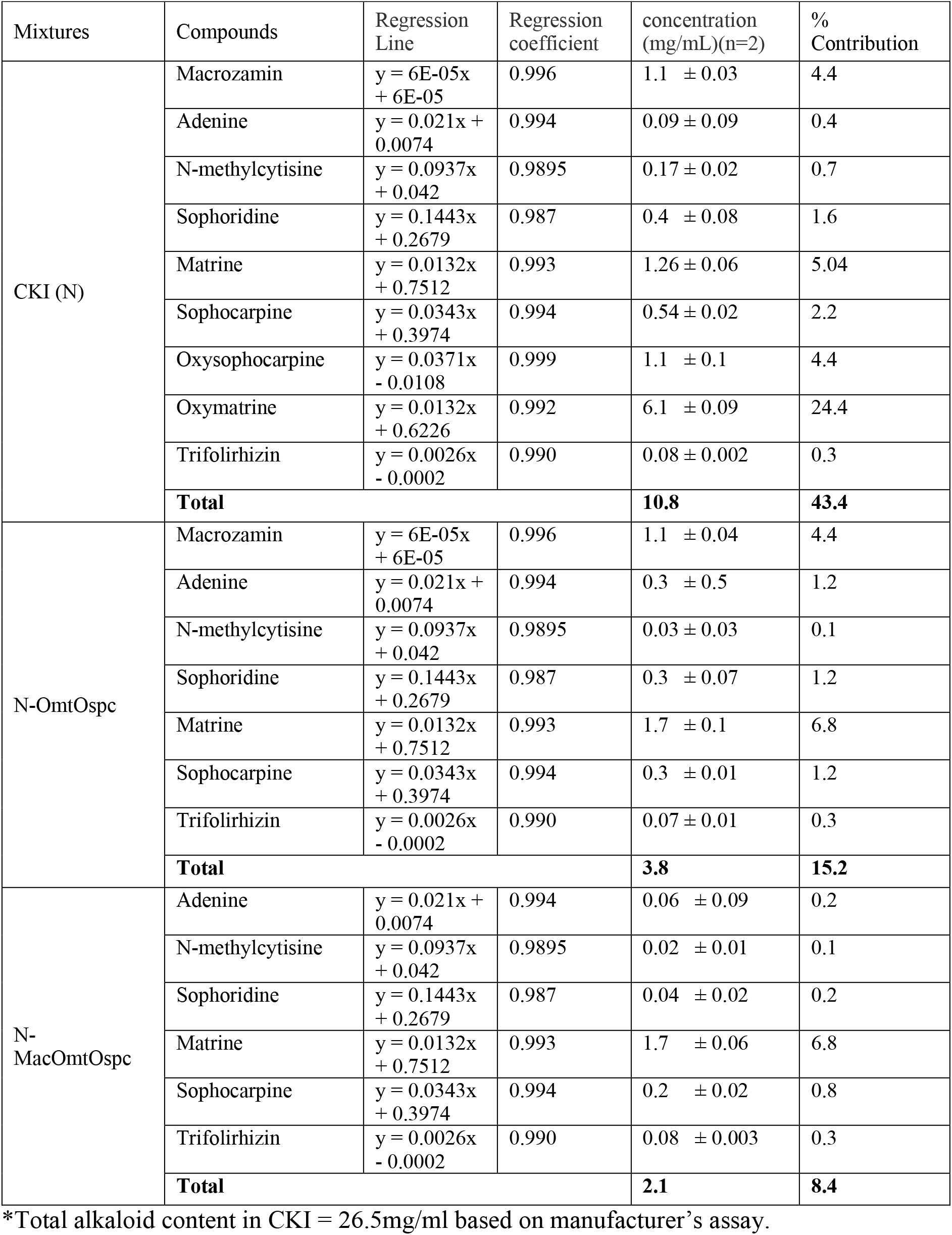
Concentration of 9 major compounds in CKI (N) (Batch No:20170322), remaining major compounds in N-OmtOspc, and remaining major compounds in N-MacOmtOspc.

| Mixtures | Compounds | Regression Line | Regression coefficient | concentration (mg/mL)(n=2) | % Contribution |
| --- | --- | --- | --- | --- | --- |
| CKI (N) | Macrozamin | $y = 6E-05x + 6E-05$ | 0.996 | $1.1 \pm 0.03$ | 4.4 |
| | Adenine | $y = 0.021x + 0.0074$ | 0.994 | $0.09 \pm 0.09$ | 0.4 |
| | N-methylcytisine | $y = 0.0937x + 0.042$ | 0.9895 | $0.17 \pm 0.02$ | 0.7 |
| | Sophoridine | $y = 0.1443x + 0.2679$ | 0.987 | $0.4 \pm 0.08$ | 1.6 |
| | Matrine | $y = 0.0132x + 0.7512$ | 0.993 | $1.26 \pm 0.06$ | 5.04 |
| | Sophocarpine | $y = 0.0343x + 0.3974$ | 0.994 | $0.54 \pm 0.02$ | 2.2 |
| | Oxysophocarpine | $y = 0.0371x - 0.0108$ | 0.999 | $1.1 \pm 0.1$ | 4.4 |
| | Oxymatrine | $y = 0.0132x + 0.6226$ | 0.992 | $6.1 \pm 0.09$ | 24.4 |
| | Trifolirhizin | $y = 0.0026x - 0.0002$ | 0.990 | $0.08 \pm 0.002$ | 0.3 |
|  | **Total** |  |  | **10.8** | **43.4** |
| N-OmtOspc | Macrozamin | $y = 6E-05x + 6E-05$ | 0.996 | $1.1 \pm 0.04$ | 4.4 |
| | Adenine | $y = 0.021x + 0.0074$ | 0.994 | $0.3 \pm 0.5$ | 1.2 |
| | N-methylcytisine | $y = 0.0937x + 0.042$ | 0.9895 | $0.03 \pm 0.03$ | 0.1 |
| | Sophoridine | $y = 0.1443x + 0.2679$ | 0.987 | $0.3 \pm 0.07$ | 1.2 |
| | Matrine | $y = 0.0132x + 0.7512$ | 0.993 | $1.7 \pm 0.1$ | 6.8 |
| | Sophocarpine | $y = 0.0343x + 0.3974$ | 0.994 | $0.3 \pm 0.01$ | 1.2 |
| | Trifolirhizin | $y = 0.0026x - 0.0002$ | 0.990 | $0.07 \pm 0.01$ | 0.3 |
|  | **Total** |  |  | **3.8** | **15.2** |
| N-MacOmtOspc | Adenine | $y = 0.021x + 0.0074$ | 0.994 | $0.06 \pm 0.09$ | 0.2 |
| | N-methylcytisine | $y = 0.0937x + 0.042$ | 0.9895 | $0.02 \pm 0.01$ | 0.1 |
| | Sophoridine | $y = 0.1443x + 0.2679$ | 0.987 | $0.04 \pm 0.02$ | 0.2 |
| | Matrine | $y = 0.0132x + 0.7512$ | 0.993 | $1.7 \pm 0.06$ | 6.8 |
| | Sophocarpine | $y = 0.0343x + 0.3974$ | 0.994 | $0.2 \pm 0.02$ | 0.8 |
| | Trifolirhizin | $y = 0.0026x - 0.0002$ | 0.990 | $0.08 \pm 0.003$ | 0.3 |
|  | **Total** |  |  | **2.1** | **8.4** |

\*Total alkaloid content in CKI = 26.5mg/ml based on manufacturer's assay.

### Phenotypic changes associated with compound deletion

Fractionation and full reconstitution caused no changes in cell viability compared to original CKI (see methods) at either 24- or 48-hours in MDA-MB-231 cells. Both reconstituted CKI and CKI caused significantly reduced viability compared to untreated (UT) cells (Supplementary Figure 2A). The MJ subtractive fraction contained a total of 9 compounds including eight previously identified MJ peaks (Ma et al., 2014) and adenine (unpublished data from Ma Yue) (Figure 1A) and the MN fraction contained the remaining peaks (Figure 1C). MJ had no effect on cell viability, while MN reduced cell viability to the same extent as CKI (Figure 2A). The 9 major compounds were individually depleted from CKI and tested as 9 (N-1) subtractive fractions, with no significant alterations in cell viability compared to CKI (Figure 2B). We then assessed the interaction effects of single MJ compounds by adding them back to the MN subtractive fraction. No change in cell viability compared to MN was observed (Supplementary Figure 2B). Sets of 3 compounds from the 9 major/standard compounds of CKI were depleted to generate 3 (N-3) subtractive fractions. The nine reference compounds were allocated into three groups, one of which contained structurally similar compounds (Omt, Ospc, Spc) and two other groups ([Mac, Ade, Tri] and [Nme, Mt, Spr]) that contained structurally different compounds. Of these three fractions, N-OmtOspcSpc decreased cell viability significantly (P<0.05) more than CKI after 48 hours (Figure 2C) while none of the sets of three compounds on their own had any effect on cell viability (Supplementary Figure 2). We then generated 9 (N-2) subtractive fractions based on the N-3 subtractive fractions (Table 1). Out of 9 (N-2) subtractive fractions (Supplementary Figure 2), only N-OmtOspc significantly decreased proliferation compared to CKI (P<0.05) (Figure 2C). We then depleted macrozamin, the only major compound derived from Baituling, together with OmtOspc as N-3 (N-MacOmtOspc) in order to determine if there was an additional effect when compared to CKI. N-OmtOspc and N-MacOmtOspc both decreased cell proliferation to the same extent (Figures 2C and 2D).

**Figure 2:**
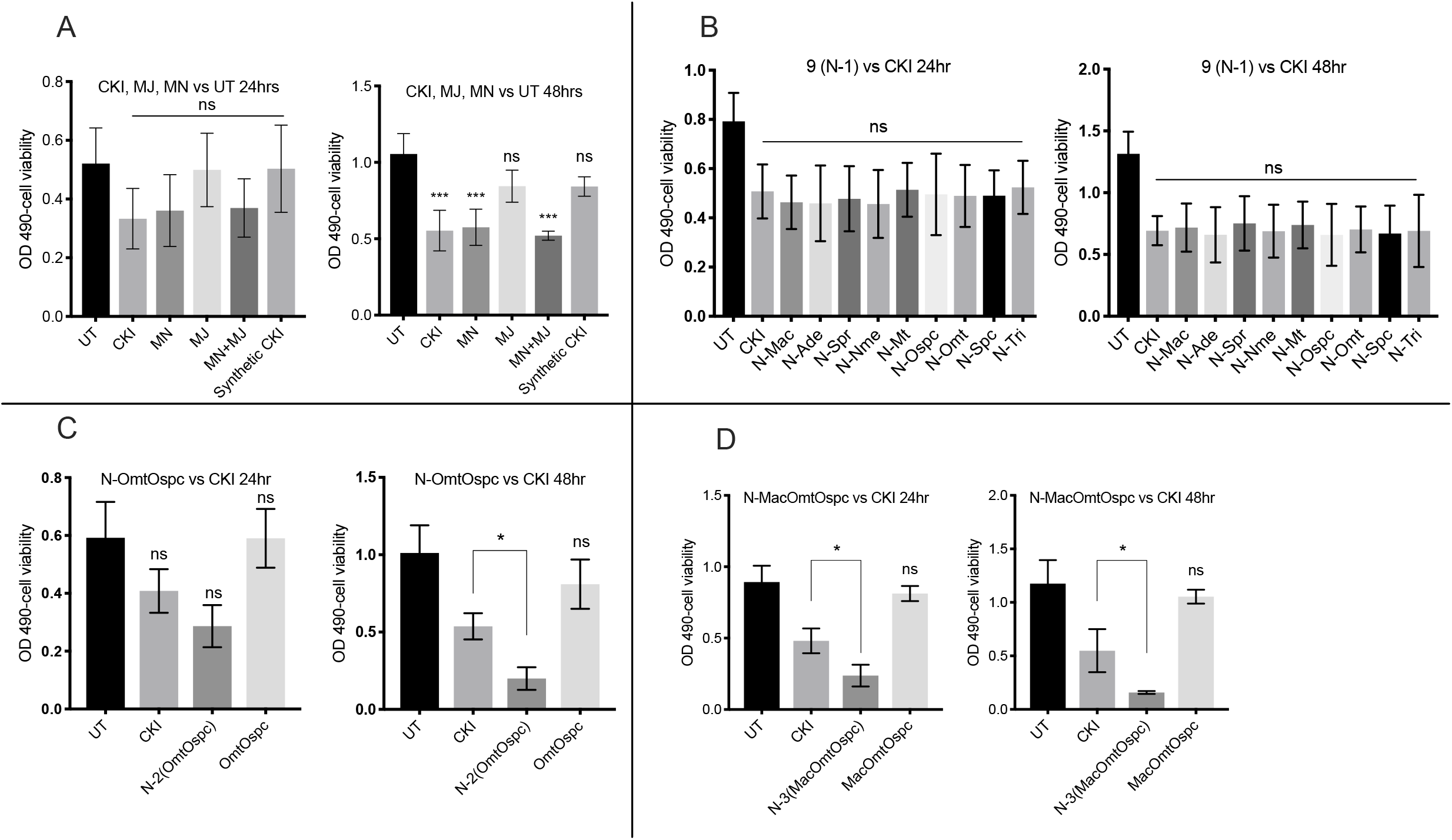
XTT Cell viability assays of subtractive fractions in MDA-MB-231 cells at 24- and 48-hour time points treated with 2mg/mL of CKI and 2mg/mL equivalent concentrations of all other treating agents. (A) Suppression of cell viability from the following fractions: UT (untreated), MJ, MN, MJ+MN (combination of MJ and MN) and Syn_CKI (synthetic CKI generated using nine major compounds). (B) Effect of 9 (N-1) subtractive fractions compared to CKI. (C) Effect of N-OmtOspc subtractive fraction and OmtOspc compared to CKI. (D) Effect of N-MacOmtOspc subtractive fraction and MacOmtOspc, compared to CKI. Statistically significant results relative to CKI treatment shown as P<0.05 (*) or ns (not significant), all data were shown as mean ± standard deviation (SD).

While no change in cell viability was found across all N-1 treatments, cell cycle analysis was performed to identify more subtle differences. There was no statistically significant difference in phases of the cell cycle of MDA-MB-231 cells for many of the N-1 treatments compared to CKI except for a statistically significant change in G1 phase by N-Omt after 48 hours (Figure 3A). On the other hand, N-OmtOspc treatment significantly altered the cell cycle for MDA-MB-231 cells and induced significant higher apoptosis from 0.25 mg/ml through 2 mg/ml treatments as compared to CKI at both time points (Figure 3B and Supplementary Figure 3). N-MacOmtOspc treatment also significantly altered the cell cycle at both time-points with generally similar effects to N-OmtOspc (Figure 3C).

**Figure 3:**
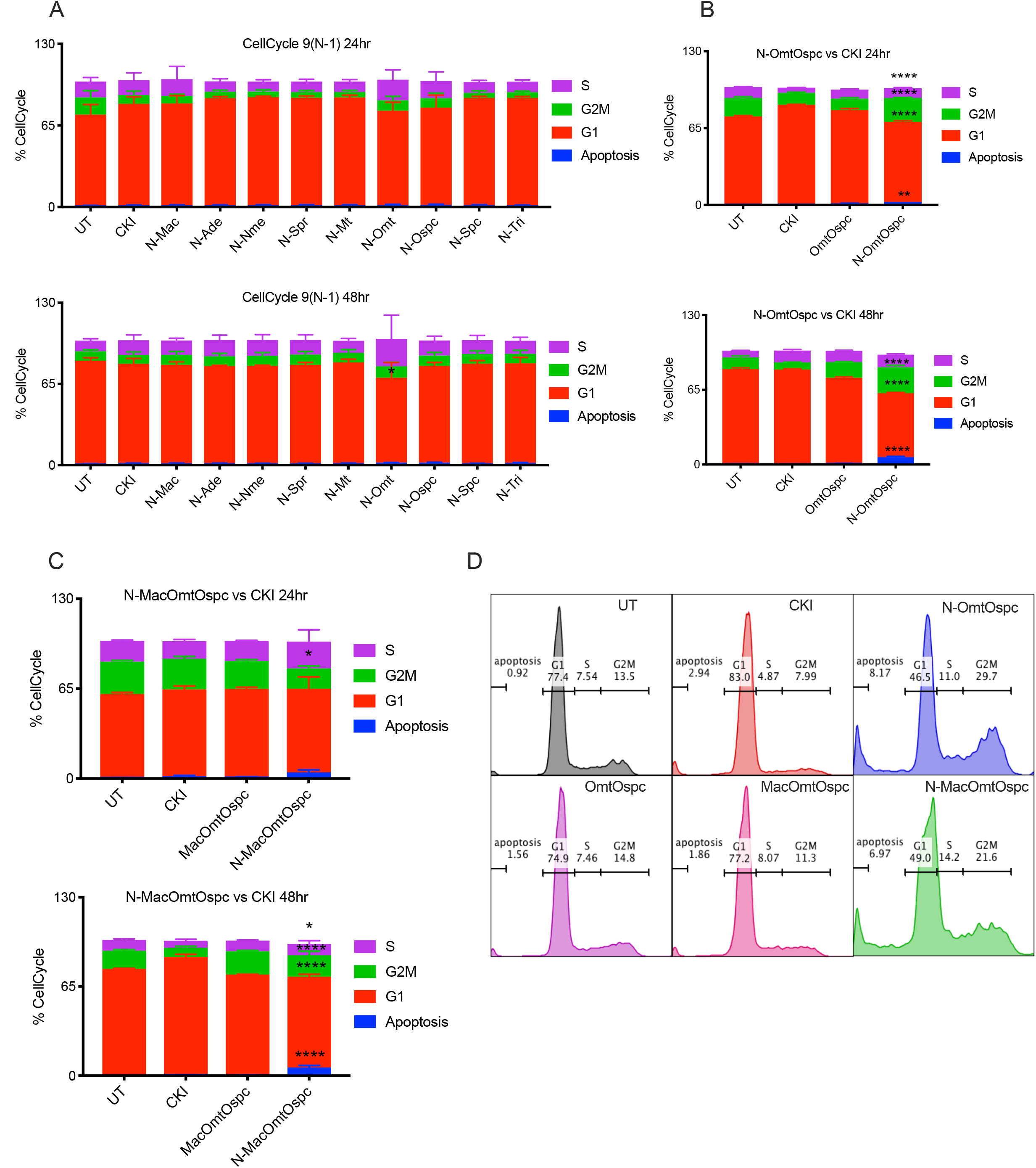
Cell cycle assay of subtractive fractions at 24- and 48-hour time point treatments. (A) Effect of 9 (N-1) subtractive fractions on cell cycle in MDA-MB-231 cells as determined by FACS PI cell cycle staining assay. (B) Effect of N-OmtOspc subtractive fraction and OmtOspc on cell cycle in MDA-MB-231 cells as determined by FACS PI cell cycle staining assay. (C) Effect of N-MacOmtOspc subtractive fraction and MOO on cell cycle in MDA-MB-231 cells as determined by FACS PI cell cycle staining assay. (D) The representative histograms of cell cycle analysis by the treatments as compared to UT. Statistically significant results shown as P<0.05 (*) or P<0.01 (**) P<0.001 (***), or P<0.0001 (****). All data were shown as mean ± standard deviation (SD).

Annexin V/PI apoptosis assays were performed using subtractive fractions on MDA-MB-231, HEK-293 (human embryonic kidney cells) and HFF (primary human foreskin fibroblasts) cell lines. While CKI at 2 mg/ml caused increased apoptosis in MDA-MB-231 cells at both 24- and 48-hour after treatment, N-OmtOspc and N-MacOmtOspc subtractive fractions at concentrations equivalent to CKI 2 mg/ml significantly increased the percentage of apoptotic cells at 24-hour with increasing apoptosis at the 48-hour timepoint, indicating that N-OmtOspc and N-MacOmtOspc significantly enhanced apoptosis compared to CKI (Figures 4A, 4E and Supplementary Figure 3A). Although CKI did not generally cause apoptosis in HEK-293 or HFF cells, N-OmtOspc and N-MacOmtOspc subtractive fractions significantly induced apoptosis (P***< 0.001) at 24-hour and 48-hour (P****< 0.0001) in both HEK-293 and HFF cells. CKI only induced apoptosis of HEK-293 (P*<0.05) at 48-hour and showed no significant apoptotic induction in HFF (Figures 4B, 4C and Supplementary Figures 3B, 3C). These results indicated that the N-OmtOspc and N-MacOmtOspc subtractive fractions induced apoptosis not only in cancerous cells but also in non-cancerous cell lines. In contrast to this, no significant apoptosis was triggered by CKI on HFF cells. A small but significant apoptotic induction was observed for HEK-293.

**Figure 4:**
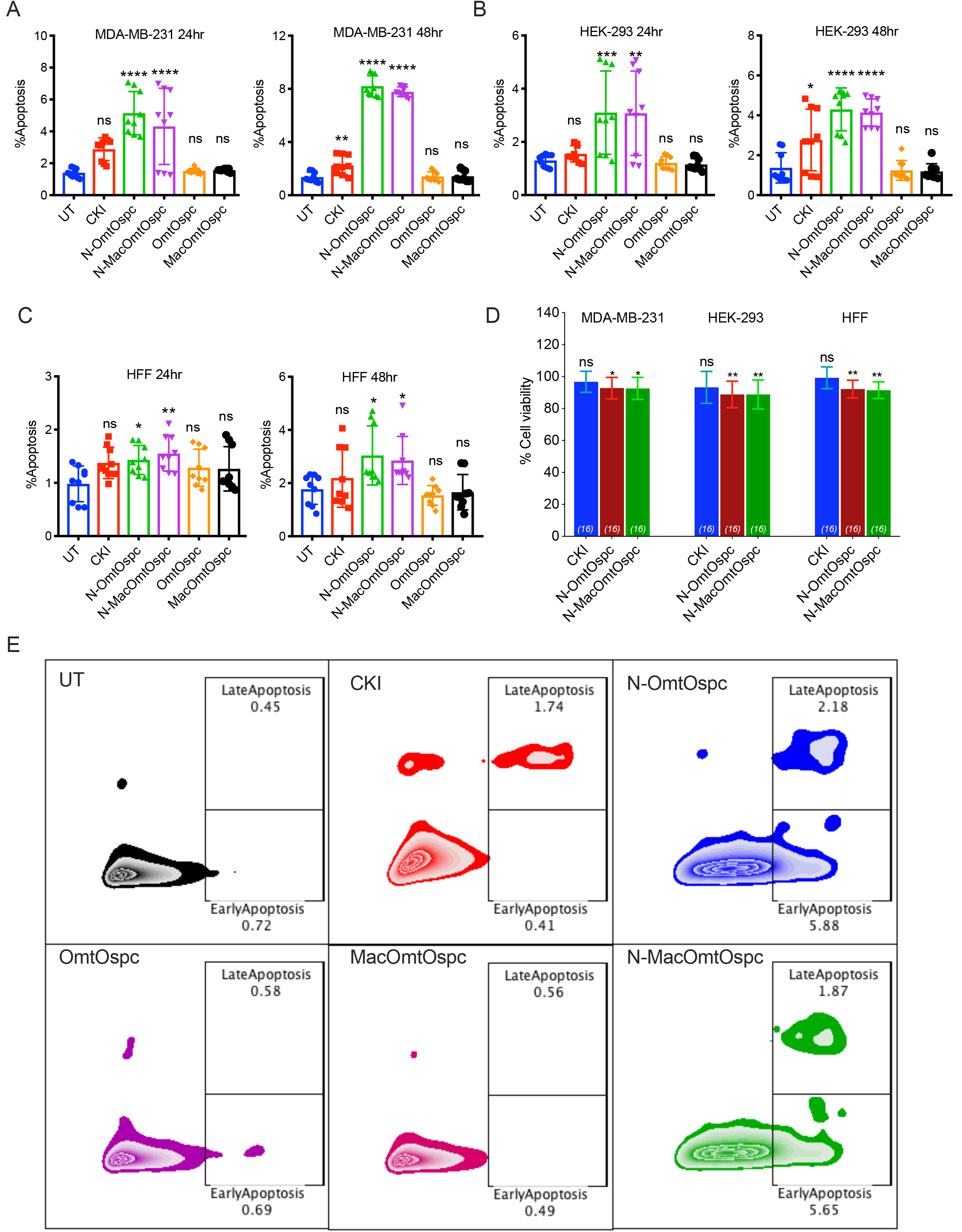
Apoptosis and cytotoxicity assays of subtractive fractions at 24- and 48-hour time point treatments. Apoptotic effect of N-OmtOspc, N-MacOmtOspc subtractive fractions, OmtOspc and MacOmtOspc in (A) MDA-MB-231 cells, (B) HEK-293 cells, and (C) HFF cells as determined by FACS AnnexinV/PI assay. (D) Cytotoxic effect of CKI, N-OmtOspc and N-MacOmtOspc subtractive fractions was determined using Alamar Blue cytotoxicity assay. Statistically significant results shown as P<0.05 (*) or P<0.01 (**) P<0.001 (***), or P<0.0001 (****); ns (not significant). All data were shown as mean ± standard deviation (SD).

Because of the significant decreased viability accompanied by increased apoptosis triggered by subtractive fractions, cytotoxicity tests were carried out for all three cell lines using CKI (2 mg/ml) and N-OmtOspc and N-MacOmtOspc subtractive fractions at concentrations equivalent to CKI 2 mg/ml. N-OmtOspc and N-MacOmtOspc at equivalent concentration to CKI 2 mg/ml were significantly cytotoxic to both non-cancerous cell lines (Figure 4D).

Overall, these results indicated that removal of combinations of specific compounds from CKI had unpredictable effects on the ability of CKI to kill cells. While removal of all major compounds from CKI caused no loss of activity and removal of all minor compounds caused total loss of activity, removal of selected major compounds (N-OmtOspc) paradoxically caused major, significant increases in the ability of CKI to reduce viability and killed cells.

### Differential gene expression

In order to understand the interactions of the components in CKI as a result of depletion, we carried out RNAseq of MDA-MB-231 cells treated with CKI and subtractive fractions. Out of nine (N-1) subtractive fractions, four, namely N-Omt, N-Mac, N-Tri and N-Nme, were selected due to their structural differences to determine their effects on transcript levels. N-OmtOspc and N-MacOmtOspc, OmtOspc, MacOmtOspc and CKI treated cells were sequenced at 24 and 48-hour time points.

After normalization, clear clustering of the replicates was observed (Figure 5A and Supplementary Figures 4, 5, 6 and 7), indicating that all 4 (N-1) treatments show comparable downstream gene expression patterns. Likewise, OmtOspc and MacOmtOspc groups and N-OmtOspc and N-MacOmtOspc groups showed similar changes in gene expression, except for one replicate (N-MacOmtOspc, 24-hour) that clustered with UT, OmtOspc and MacOmtOspc. The number of differentially expressed (DE) genes associated with each treatment was calculated using pair-wise comparative analysis. CKI treatment was used as a baseline to compare all other treatments in order to emphasize the effect of depleted compounds and CKI treatment was compared to UT.

**Figure 5:**
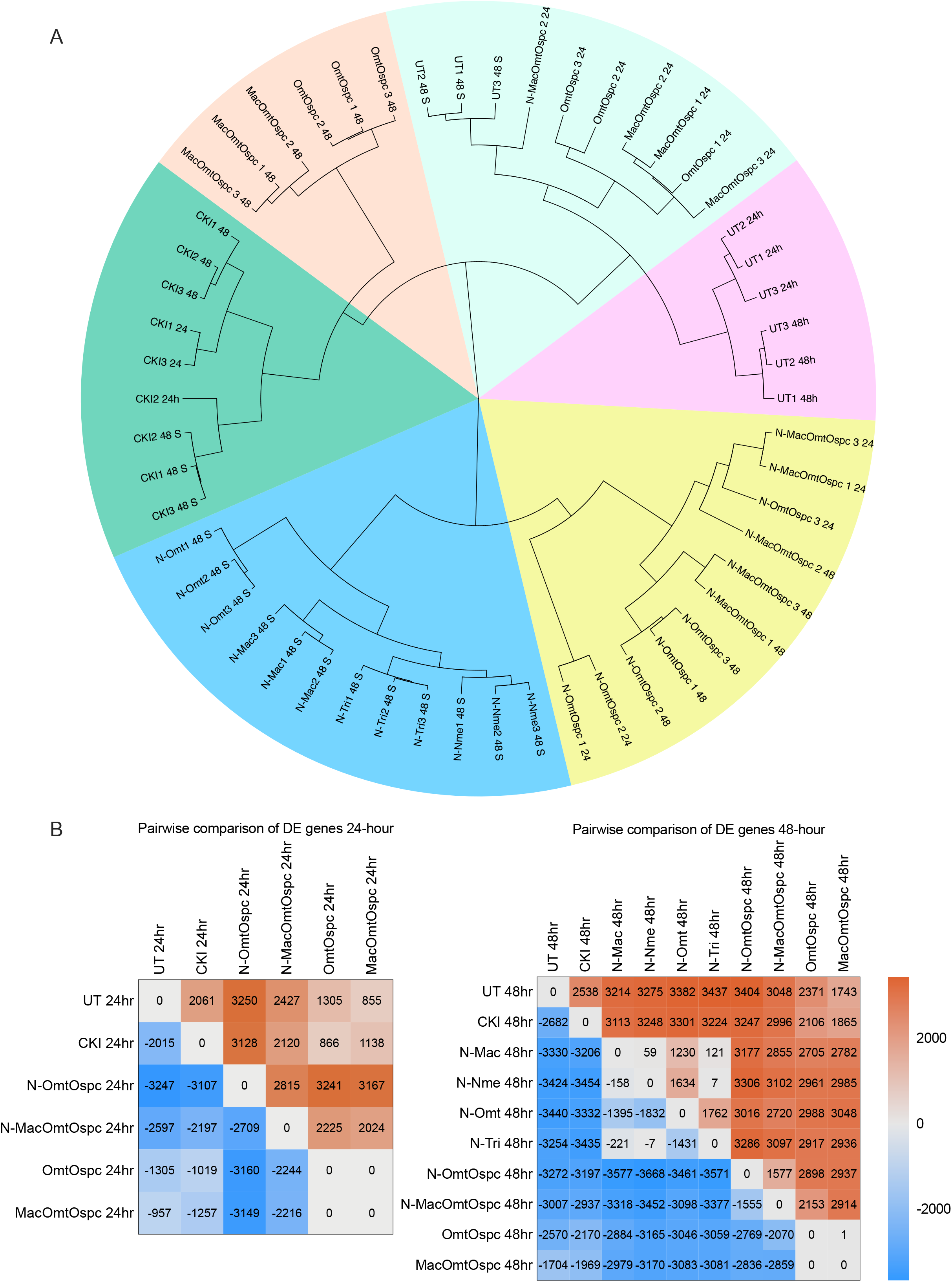
Gene expression clustering and summary of differential gene expression (A) Clustering of treated samples based on gene expression, calculated as transcripts per million (TPM) using Ward’s hierarchical cluster analysis (Ward.D2) method. Number of differentially expressed (DE) genes (FDR < 0.05 according to edgeR) associated with each treatment was calculated using pair-wise comparison at (B) 24 hours and (C) 48 hours time point. Treatments were compared column versus row. Up-regulated genes are shown in shades of red and down-regulated genes are shown in shades of blue.

There were thousands of upregulated and downregulated genes at 24 and 48 hours in most pairwise comparisons (Figure 5B). However DE genes between OmtOspc and MacOmtOspc treatments were not observed and there were almost no DE genes between N-Mac, N-Nme and N-Tri treatments (Figure 5B) indicating that these three subtractive fractions had very similar effects on gene expression.

When we compared the DE genes found between treatments, there were a large number of DE genes (~71.3%) shared between all four (N-1) treatments (Supplementary Figures 8 and 9 and Supplementary Table 3). A similar number of shared DE genes (~24.6%) between four (N-1), OmtOspc and MacOmtOspc and between four (N-1), N-OmtOspc and N-MacOmtOspc as compared to CKI at 48-hours indicated that gene expression patterns from N-1 treatments were mostly different from N-OmtOspc, N-MacOmtOspc, OmtOspc and MacOmtOspc treated cells. 55% of the DE genes between UT, OmtOspc and MacOmtOspc were shared. When the four (N-1) treatments were compared to CKI treatment, 42.8% of DE genes were shared, and when N-OmtOspc and N-MacOmtOspc treatments were compared to CKI, 50.1% DE genes were shared, indicating that N-OmtOspc and N-MacOmtOspc treatments appeared to be more similar to CKI than N-1 treatments.

**Table 3:** Summary of shared, differentially expressed (DE) genes across treatments. Similarity (%) calculated from total number of shared DE genes from all listed comparisons. To find the number of DE genes, CKI treatment was used as a baseline to compare all other fractionated treatments in order to emphasize the effect of depleted compounds and UT (untreated) was used as a base to calculate the DE genes for CKI treatment.

| Treatments Comparisons | Similarity (%) | Time course (hours) |
| --- | --- | --- |
| 4 x (N-1) | 71.3 | 48 |
| 4 x (N-1) and CKI | 42.7 | 48 |
| 4 x (N-1), CKI and N-OmtOspc | 30.4 | 48 |
| 4 x (N-1), CKI, N-OmtOspc and N-MacOmtOspc | 24.6 | 48 |
| 4 x (N-1), CKI, OmtOspc, and MacOmtOspc | 24.6 | 48 |
| 4 x (N-1), OmtOspc, and MacOmtOspc | 33.7 | 48 |
| 4 x (N-1), N-OmtOspc and N-MacOmtOspc | 37.7 | 48 |
| CKI and N-OmtOspc | 64.4 | 48 |
| CKI and N-MacOmtOspc | 63.9 | 48 |
| CKI, N-OmtOspc and N-MacOmtOspc | 50.1 | 48 |
| CKI, N-OmtOspc, N-MacOmtOspc, OmtOspc, and MacOmtOspc | 30.2 | 48 |
| UT, OmtOspc, and MacOmtOspc | 54.9 | 48 |
| CKI and N-OmtOspc | 56.0 | 24 |
| CKI and N-MacOmtOspc | 45.6 | 24 |
| CKI, N-OmtOspc and N-MacOmtOspc | 31.6 | 24 |
| CKI, N-OmtOspc, N-MacOmtOspc, OmtOspc, and MacOmtOspc | 13.3 | 24 |
| UT, OmtOspc, and MacOmtOspc | 39.1 | 24 |

The overall levels of similarity in DE genes were as follows: 1)All N-1 treatments had approximately 70 % similar gene expression patterns, 2) OmtOspc and MacOmtOspc treatments were approximately 50% similar to UT and 33% similar to N-1 treatments, 3) downstream gene expression patterns between N-1, N-OmtOspc and N-MacOmtOspc were approximately 37% similar.

### Gene ontology and pathway annotation of DE genes

DE genes were analysed for over-representation in our data sets with respect to biological function using Gene Ontology (GO) annotation. We looked for shared DE genes between treatments and identified over-represented genes in these shared genes. The only common function enriched across all comparisons was for "cell cycle checkpoint" (Figure 6A). This confirmed earlier results (Qu et al., 2016) and was consistent with the phenotype data for CKI.

**Figure 6:**
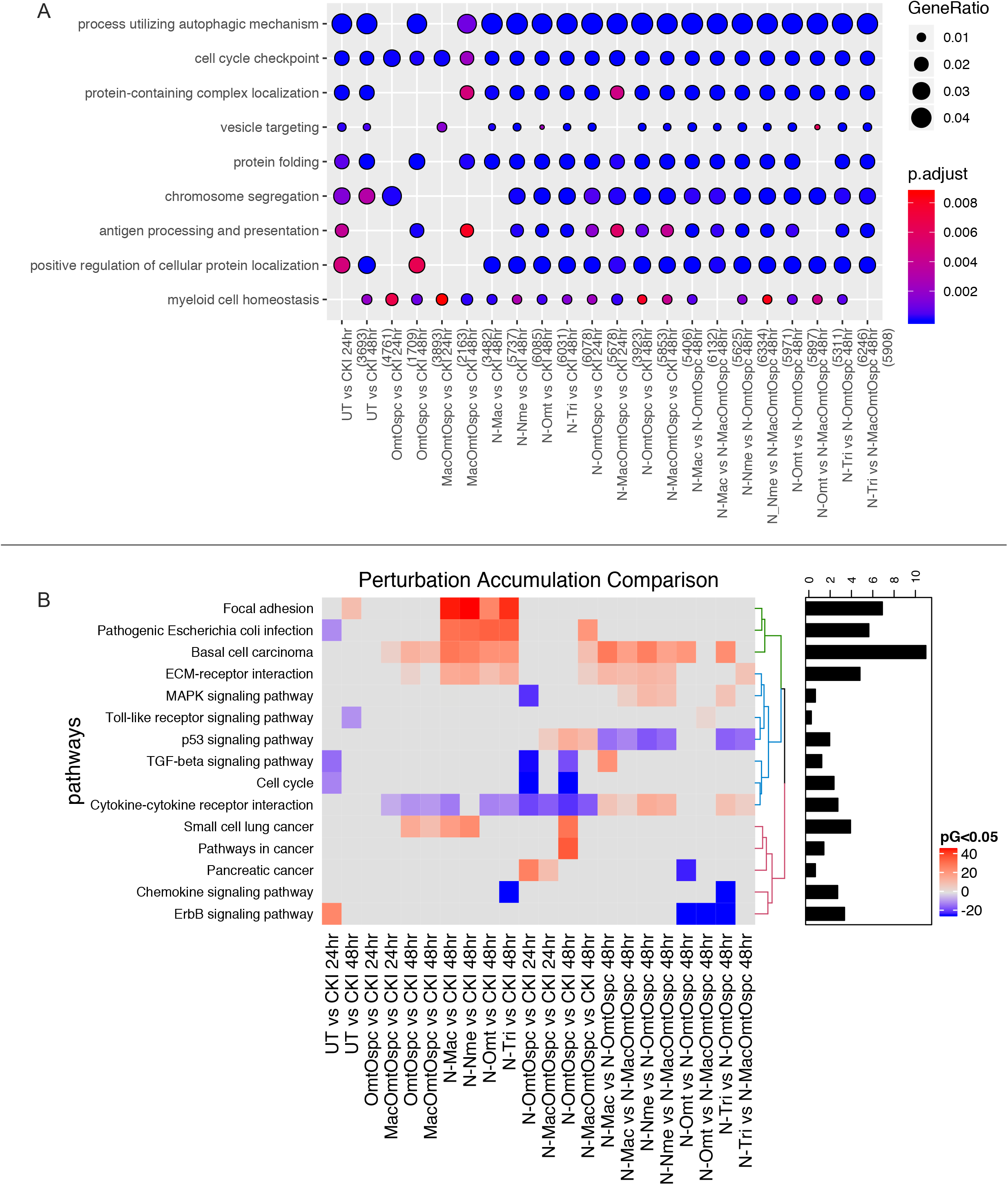
Over-representation analysis of GO functional annotation and KEGG pathway perturbation analysis. (A) Over-represented GO terms (Biological Process, BP=3) for DE genes identified from comparison of subtractive fraction treated cells against CKI treatment in order to show the relative change from depleting compounds. Gene ratio of each term calculated from Cluster Profiler was plotted based on the adjusted *P*-values. Top 5 most significant categories of GO terms were plotted by default. Colour gradient of adjusted *P*-values ranging from red to blue in order of increasing p-values (high to low significance). Number of identified genes in each treatment (numbers in parentheses) were shown in the bottom and the sizes of the dots correspond to the ratio of gens out of all significant DE gene from each treatment involved in the particular terms. (B) Identification of significantly perturbed pathways using SPIA (pG < 0.05) analysis. Eighty-six significantly perturbed pathways from twenty-two comparisons were found (Supplementary Figure10). Only 15 pathways most obviously linked to our phenotypes of cell viability, cell cycle and apoptosis were shown here. Positive (overall increase in gene expression for pathway) and negative (overall decrease in gene expression for pathway) perturbation accumulation values of the pathways were shown in red and blue respectively. Mean perturbation values of each pathway were shown in bar plot.

### Subtracted fractionation altered pathways

We also performed pathway based analysis to look for pathway level perturbation by comparing DE genes within Kyoto Encyclopedia of Genes and Genomes (KEGG) pathways between treatments. We used Signaling Pathway Impact Analysis (SPIA) to identify pathways with statistically significant perturbation values expected to alter pathway flux. We identified 86 pathways (Supplementary Figure 11) with statistically significant (P< 0.05) perturbations of gene expression and of these, 15 pathways were most obviously linked to our phenotypes of cell viability, cell cycle and apoptosis (Figure 6B). By comparing the pathway gene expression perturbation scores (pG) between treatments three specific observations could be made: 1) N-1 fractional deletions vs CKI had significant effects on flux in some pathways without phenotypic effects, 2) N–OmtOspc vs CKI which had a pronounced phenotypic effect at both 24- and 48-hours, had a significant effect on reducing estimated pathway flux for Cytokine-Cytokine Receptor, Cell Cycle and TGF-Beta signaling pathways, 3) comparison of N-1 fractional deletions vs fractional deletions of N–OmtOspc/N–MacOmtOspc showed consistent pathway perturbations for Cytokine-Cytokine Receptor and p53 signaling pathways. On this basis, we inferred that different major compounds could be deleted with very similar effects, indicating that they may have similar targets. In contrast, deleting Omt and Ospc simultaneously caused a significant shift in phenotype and was accompanied by specific perturbations in pathways that regulate inflammation, cell cycle and apoptosis. The combined deletion of Omt and Ospc had a synergistic effect on viability, cell-cycle and apoptosis and a synergistic effect on gene expression, consistent with the observed changes in pathway specific perturbation of gene expression. Because this double compound deletion potentiated the cell killing effect of CKI we hypothesised that the compounds in CKI have multiple targets leading to a phenotypic effect that reflects the integration of stimulation and inhibition across all those targets. Removal of Omt and Ospc alter the balance of stimulation and inhibition leading to an integrated effect for the remaining compounds in the mixture that caused more cell death than CKI.

More detailed examination of some of these interactions within significantly perturbed pathways highlighted the gene-specific changes in expression for some key regulators of inflammation and the cell-cycle. Most effects on gene expression from deletion of single vs two compounds were similar, suggesting that the enhanced cell killing by N-OmtOspc was due to additive effects of the compound deletions. However, by comparing differences in pairwise comparisons between treatments at the gene level within the Cytokine-Cytokine Receptor Interaction and Cell Cycle pathways we identified a subset of genes that had opposite changes in gene expression when comparing single compound deletions to N-OmtOspc deletion. In the Cytokine-Cytokine Receptor Interaction pathway (Figure 7) these genes are IL1-R1 (Interleukin-1 Receptor), IL-27RA (Interleukin-27 Receptor alphasubunit), TNFRSF1B (Tumor Necrosis Factor Receptor Superfamily Member 1B), TNFRSF14 (Tumor Necrosis Factor Receptor Superfamily, Member 14) and OSMR (Oncostatin M Receptor/IL-31 Receptor Subunit Beta) and they all transduce inflammatory ligand signals to the NFkB pathway and/or the apoptosis pathway. In the Cell-Cycle pathway (Figure 8) these genes are CDKN1C (Cyclin-Dependent Kinase Inhibitor 1C (P57, Kip2), CDC25B (Cell Division Cycle 25B), ATR (ATR Serine/Threonine Kinase), CDKN1B (Cyclin-Dependent Kinase Inhibitor 1B (P27, Kip1)), CDKN2D (Cyclin-Dependent Kinase Inhibitor 2D (P19, Inhibits CDK4)), TGFB1 (Transforming Growth Factor Beta 1), FZR1 (Fizzy And Cell Division Cycle 20 Related 1), CDC20 (Cell Division Cycle 20), CDC27 (Cell Division Cycle 27), ORC2 (Origin Recognition Complex Subunit 2), ANAPC4 (Anaphase Promoting Complex Subunit 4), ZBTB17 (Zinc Finger And BTB Domain Containing 17) and ABL1 (ABL Proto-Oncogene 1, Non-Receptor Tyrosine Kinase). The opposite changes in gene expression stimulated by N-OmtOspc compared to N-1 subfractions provides support for the idea that multiple major compounds can have similar effects on specific genes but that the combination of Omt and Ospc can have synergistic and opposite effects on those same genes. This means that multiple compounds with overlapping targets (based on their structural similarities) can either reinforce a single outcome or exhibit unpredictable and opposite effects when combined.

**Figure 7:**
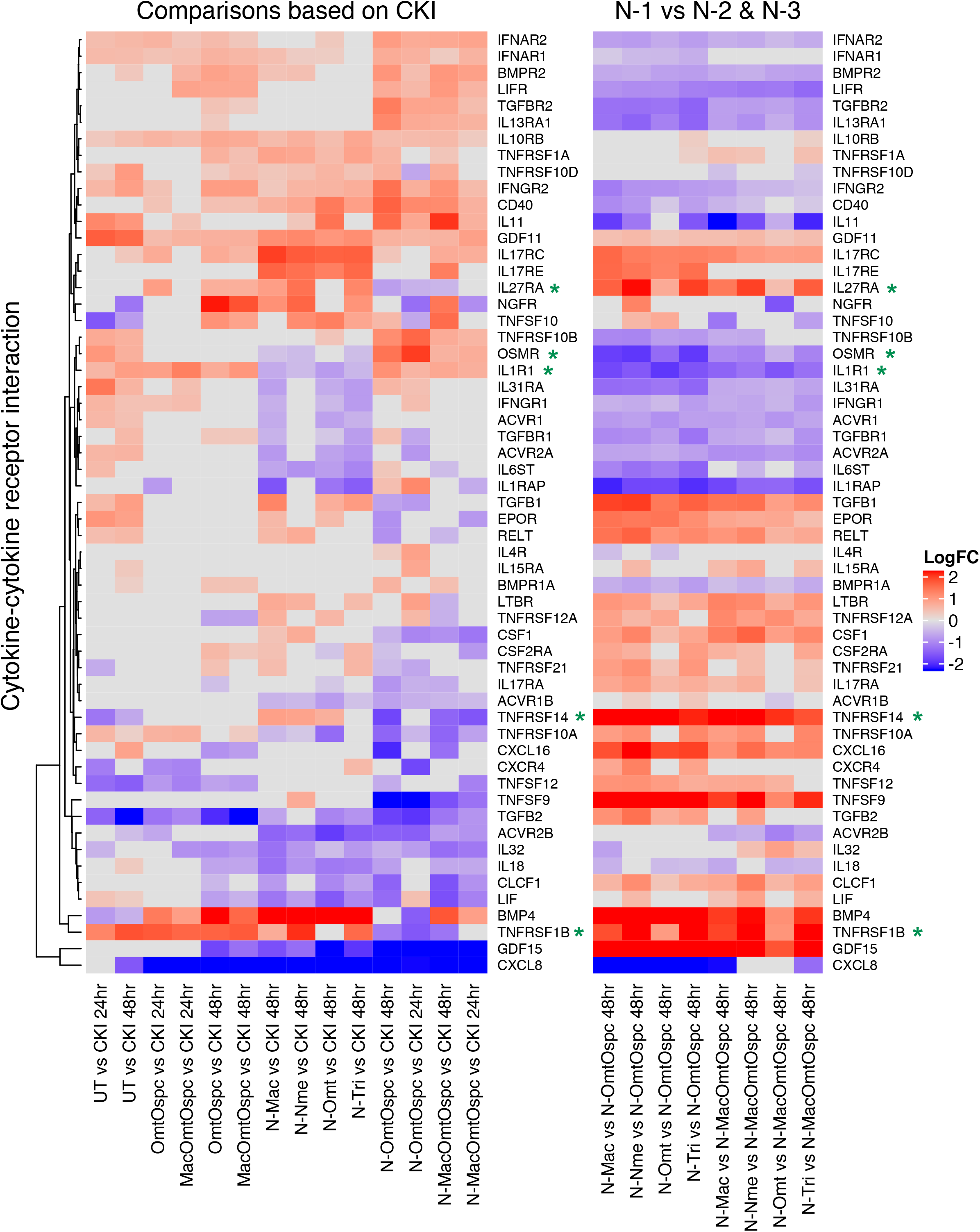
Differential gene expression profiles of all treatments for Cytokine-Cytokine Receptor pathway. The left panel shows comparison of subtractive fraction treated cells against CKI treatment and the right panel shows comparison of single compound subtractive fraction treated cells against the treatments for two and three compound subtractive fractions. Asterisks in green shows a subset of genes that had opposite changes in gene expression across treatments.

**Figure 8:**
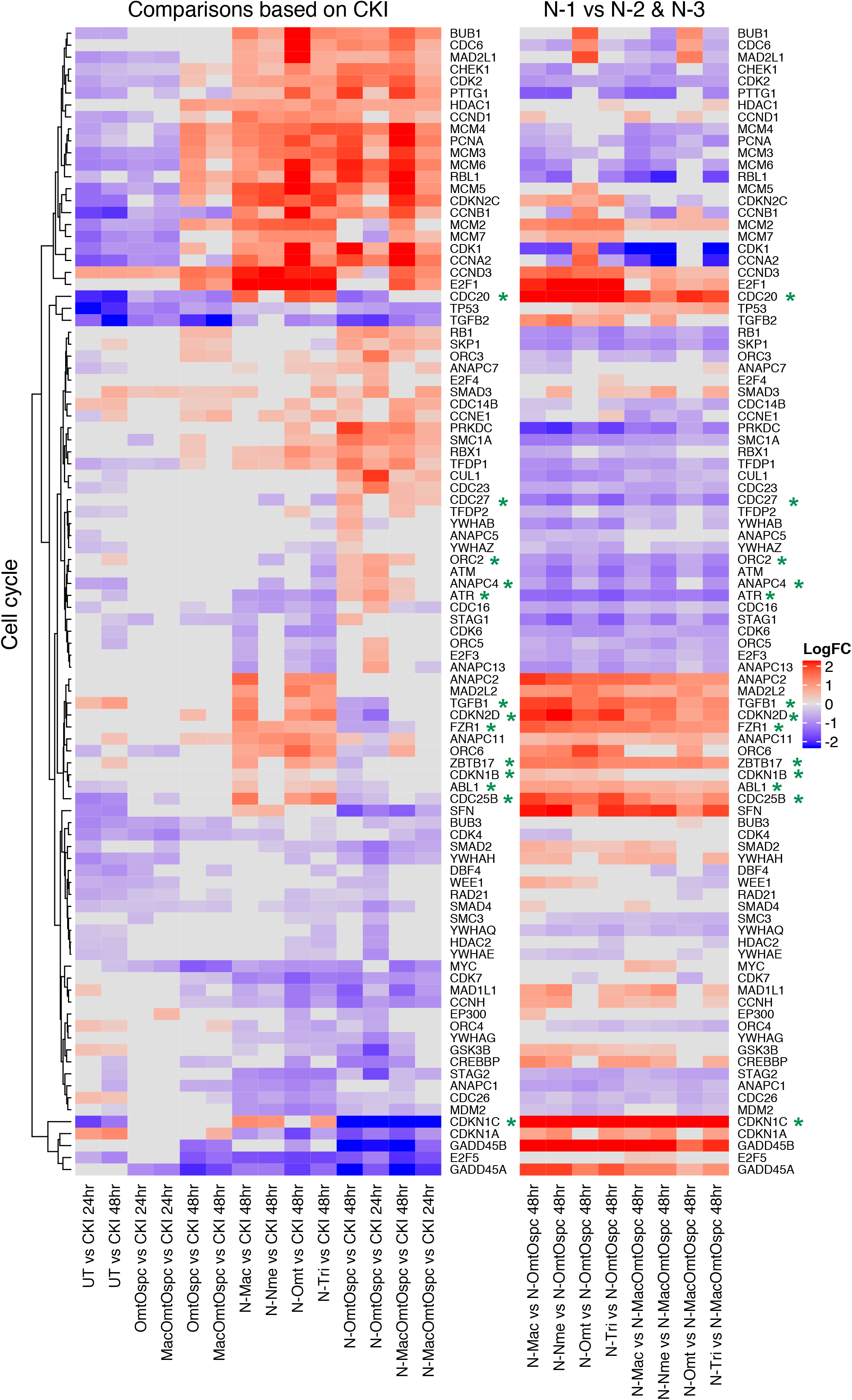
Differential gene expression profiles of all treatments for Cell Cycle pathway. The left panel shows comparison of subtractive fraction treated cells against CKI treatment and the right panel shows comparison of single compound subtractive fraction treated cells against the treatments for two and three compound subtractive fractions. Asterisks in green shows a subset of genes that have opposite changes in gene expression across treatments.

Overall our results support the concept of multi-compound/multi-target interactions for plant extract based drugs that contain many plant secondary metabolites. Biological effects of complex plant extracts may result from interactions of multiple compounds, with negligible effects from single compounds alone. This has implications for how we assess the functional evidence for such extracts.

## Discussion

Previous studies have demonstrated that CKI can alter the cell cycle, induce apoptosis and reduce proliferation in various cancer cell lines (Xu et al., 2011;Qu et al., 2016;Gao et al., 2018). CKI also killed leukaemia cells via the Prdxs/ROS/Trx1 signalling pathway in an acute myeloid leukaemia patient-derived xenograft model and caused cell cycle arrest in U937 leukaemia derived cells (Jin et al., 2018). Cell cycle arrest by CKI at checkpoints is correlated with the induction of double strand breaks by CKI treatment (Cui et al., 2018). In contrast to our experiments reported above, oxymatrine was previously shown to arrest the cell cycle and induce apoptosis in human glioblastoma cells through EGFR/PI3K/Akt/mTOR signaling pathway (Dai et al., 2018) and inhibit the proliferation of laryngeal squamous cell carcinoma Hep-2 cells (Ying et al., 2015). As shown in this report, oxymatrine or oxysophocarpine or combined OmtOspc treatment caused no significant change in cell viability, the cell cycle or apoptosis, in agreement with prior work that showed oxymatrine and oxysophocarpine exerting no significant effect on apoptosis, cell cycle or cell proliferation in HCT116 human colon cancer cells (Zhang et al., 2014).

The paradoxical result that removal of OmtOspc caused a striking increase in apoptosis is most simply explained by a model based on integrating effects of multiple compounds on many targets. The interactions between compounds in the mixture can be synergistic and antagonistic such that if two compounds are removed that have a synergistic effect that is antagonistic to the remainder of the mixture, the resulting depleted mixture will be dis-inhibited compared to CKI. This is illustrated by our studies and others that show single compounds alone had no or little effect compared to CKI. For instance, while CKI treatment resulted in increased DNA double strand breaks and affected the cell cycle resulting in decreased cancer cell proliferation, oxymatrine alone exhibited only a small effect in the same assay (Cui et al., 2018). Gao and colleagues also reported that oxysophocarpine at 4 mg/ml had no effect, oxymatrine at 4 mg/ml (*P<0.05) and CKI at 2 mg/ml (***P<0.001) significantly reduced the proliferation of hepatocellular carcinoma SMMC-7721 cells *in vitro* (Gao et al., 2018). Although significant inhibition of proliferation by oxymatrine occurred, the concentration used in this experiment was ~ 8x times higher than that of oxymatrine in 2 mg/ml of CKI. These studies agree with our experimental outcomes that oxymatrine and oxysophocarpine individually had no or little effect compared to CKI treatment.

At the level of gene expression in our study, gene ontology analysis indicated that genes for “cell cycle checkpoint” were significantly enriched in cells treated with all fractionated mixtures or mixtures of Omt and Ospc. Consistent with other studies, our results also demonstrated that these compounds had little or no phenotypic effect on their own, but that when both were deleted, the remaining compounds unexpectedly had significantly greater effects on phenotype and gene expression. When examined in the context of specific pathways, treatment with OmtOspc or N-OmtOspc which had strikingly different effects on phenotype, had similar effects on the perturbation of the “Cytokine-Cytokine Receptor Interaction” pathway, the most commonly perturbed pathway seen in our analysis, that interestingly did not show up when comparing CKI to UT. This is consistent with previous information showing that CKI induced cytokines IL4 and IL10 in cancer patients with acute leukaemia (Tu et al., 2016). In contrast to this observation, IL4 and IL10 levels were significantly decreased in transgenic mice treated with oxymatrine at a dose of 200 mg/kg (Dong et al., 2002). In our experiment, we also observed that while CKI and many of the depleted fractions had significant effect on the genes in the “Cytokine-Cytokine Receptor Interaction” pathway, OmtOspc and MacOmtOspc had little effect on the genes in that pathway. The observation that many genes in the “Cytokine-Cytokine Receptor Interaction” pathway were not affected by OmtOspc and MacOmtOspc compared to deletion fractions confirmed that removal of compounds rather than treatment with single or a few compounds can be more informative of the role and significance of individual compounds as part of mixtures/extracts.

In summary, Our approach allowed the identification of both synergistic and antagonistic interactions within the drug mixture. Viewed as a network where the compounds and the targets are nodes and the interactions between compounds and targets, and between targets are edges, it is clear that the edges (interactions) determine the overall effect of the compound mixture. By removing one or two compounds from a mixture, we can potentially perturb the target network(s) to either reduce the effect of the mixture for some outcome or potentiate it for another. We believe this approach may be of general use for the study of herbal medicines/extracts, avoiding failures that stem from exclusive reliance on the identification of a single compound that accounts for most of the biological activity in mixtures.

## Materials and Methods

### Cell lines

MDA-MB-231 cells were purchased from the American Type Culture Collection (ATCC, Manassas, VA). HEK-293 and HFF were kindly provided by Prof. Andrea Yool (Medical School, University of Adelaide). Cells were cultured in DMEM (Dulbecco’s Modified Eagle’s Medium, Invitrogen) with 10% FBS (Fetal bovine serum, Thermo Fisher Scientific) at 370C with 5% CO_2_.

### Compound fractionation by HPLC

CKI (Batch No: 20170322, total alkaloid concentration of 26.5 mg/ml) was provided by Zhendong Pharmaceutical Co.Ltd (Shanxi, China). CKI (N) was processed to deplete Single (N-1), double (N-2) and triple (N-3) compounds using HPLC by standardizing using nine compounds, namely Oxymatrine (Omt), Oxysophocarpine (Ospc), N-Methylcytisine (Nme), Matrine (Mt), Sophocarpine (Spc), Trifolirhizin (Tri), Adenine (Ade), Sophoridine (Spr) (Beina Biotechnology Institute Co., Ltd, China), and macrozamin (Zhendong Pharmaceutical Co.Ltd) which were previously reported to be found in published and unpublished data (Ma et al., 2014). HPLC fractionation separated Minor (MN) and Major (MJ) peaks to determine the principle and secondary components. The MJ mixture contained the nine standard compounds mentioned above and MN contained the remaining CKI components. In addition, nine N-1 fractional deletions, nine N-2 fractional deletions and three N-3 fractional deletions were produced.

HPLC separation was achieved using an HPLC instrument (Shimadzu, Japan) equipped with a photodiode-array UV-Vis detector with preparative C_18_ column (5 μm, 250 × 10 mm) (Phenomenex, Australia). The following mobile phase was used to fractionate the CKI mixture: 0.01 M ammonium acetate (adjusted to pH 8.0, solvent A) and acetonitrile + 0.09 % trifluroacetic acid (solvent B). The flow rate was 2 ml/min and linear gradient was adopted as follows; 0 min, 100% A; 60 min, 65% A, 70 min, 100%A. The chromatogram was recorded from 200 nm to 280 nm, with monitoring at 215 nm. Samples were frozen and lyophilised using a Christ Alpha 1-2 LD lyophilizer (Martin Christ Gefriertrocknungsanlagen GmbH, Germany). Several cycles of lyophilisation and resuspension were used to remove all remaining HPLC solvents and final reconstitution was carried out using MilliQ water buffered with 10 mM Hepes (Gibco, Life technologies, USA) and adjusted to pH 6.8-7.0. Lyophilised samples were resuspended to create an equivalent dilution for compounds in the sample compared to CKI.

### Identification of reconstituted mixtures by LC-MS/MS

Agilent 6230 TOF mass spectrometer was used to determine the concentration of the known compounds from the CKI and reconstituted N-OmtOspc and N-MacOmtOspc mixtures. 10uL sample was injected with the flow rate of 0.8 ml/min, a gradient program of 0 min, 100 % A; 40 % B; 25 min, 60 % B, 35 min, and solvents H_2_O + 0.1 % formic acid (solvent A) and acetonitrile + 0.1% formic acid (solvent B). The column used was C18 (5μ, 150 x 4.6 mm, Diamosnsil, Dkimatech). The recovered contents of the samples was measured by spike-in compound cytosine. Gas phase ions were generated with an electrospray source, with with key instrument parameters: gas temperature, 325; sheath gas temperature, 350; vCap, 3500; fragmentor, 175; acquisition range (m/z) 60-17000. Calibration curves for 9 standard compounds containing various concentrations were shown in Supplementary Data 1.

### Cell viability Assay

XTT [2,3-bis-(2-methoxy-4-nitro-5-sulfophenyl)-2*H*-tetrazolium-5-carboxanilide] and PMS (N-methyl dibenzopyrazine methyl sulfate) (50:1, Sigma-Aldrich) assay was used to assess cell viability following. Briefly, 8,000 cells in 50 ul of medium were plated in 96 wells trays overnight prior to drug treatments in triplicate. Cells were subsequently treated with 50ul of drug mixtures to provide final concentrations of 0.25, 0.5, 2 and 2 mg/ml in 100 μl. Cell viability was then measured at 24-and 48-hours after drug treatment by the addition of 50 μl of XTT:PMS (50:1 ratio). An equal volume of medium and treating agents plus XTT:PMS was used to subtract the background optical density (OD). The absorbance of each well was recorded using a Biotrack II microplate reader at 492 nm.

### Annexin V/PI apoptosis assay

Apoptosis, or programmed cell death, resulting from treatment was determined using an Annexin V-FITC apoptosis detection kit (eBioscience™ Annexin V-FITC Apoptosis Detection Kit, Thermofisher Scientific) according to the manufacturer’s protocol. Briefly, 4×105 cells were seeded in 6-well plates in triplicate overnight prior to treatment. On the following day, cells were treated with the agents as described for 24-and 48-hours. Data were acquired with a BD LSR-FORTESSA (NJ, USA) flow cytometer, and FlowJo software (TreeStar Inc., OR, USA) was used to analyse the acquired data and produce percent apoptosis values.

### Cell cycle assay

Cell culture and drug treatments were performed as described above for cell cycle analysis. A Propidium Iodide (PI) staining protocol (Riccardi and Nicoletti, 2006) was used to detect the changes in cell cycle as a result of treatment after 24- and 48-hours. The characteristics of stained cells were measured using a BD LSR-FORTESSA (NJ, USA) flow cytometer, and acquired data were analysed using FlowJo software.

### Cytotoxicity assay

MDA-MB-231, HEK-239 and HFF cells were seeded in 96-well plates at a density of 2.5 × 103 cells per well in triplicate. CKI and fractionated mixtures to produce a final concentration of 1 mg/ml and 2 mg/ml were added to each well and after 24-hours of incubation and viable cells were measured using the Alamar Blue assay (Molecular Probes, Eugene, OR). Mercuric chloride (Sigma-Aldrich, St. Louis, MO) (5 μM) was used as a positive control and wells without cells were set as a negative control in the same plate.

### Sample preparation and RNA sequencing

Cells were plated in 6 well plates with a density of 2×105 cells/ml overnight prior to drug treatments. On the following day, CKI (to give a final concentration of 2 mg/ml) and fractionated mixtures (equivalent dilutions of CKI) were added. Total RNA was isolated by using an RNA extraction kit (Thermo Fisher Scientific) according to the manufacturer’s instructions and RNA samples were quantified and quality determined using a Bioanalyzer at the Cancer Genome Facility of the Australian Cancer Research Foundatin (SA, Australia). RNA samples with RNA integrity number (RINs) > 7.0 were sent to be sequenced at Novogene (China). Briefly, after QC procedures were performed, mRNA was isolated using oligo(dT) beads and randomly fragmented by adding fragmentation buffer, followed by cDNA synthesis primed with random hexamers. Next, a custom second-strand synthesis buffer (Illumina), dNTPs, RNase H and DNA polymerase I were added for second-strand synthesis After end repair, barcode ligation and sequencing adaptor ligation, the double-stranded cDNA library was size selected and PCR amplified. Sequencing was carried out on an Illumina HiSeq X platform with paired-end 150 bp reads.

### Transcriptome data processing

FastQC (v0.11.4, Babraham Bioinformatics) was used to check the quality of raw reads before proceeding with downstream analysis. Trim_galore (v0.3.7, Babraham Bioinformatics) with the parameters: --stringency 5 --paired --fastqc_args was used to trim adaptors and low-quality sequences. STAR (v2.5.3a) was then applied to align the trimmed reads to the reference genome (hg19, UCSC) with the parameters: --outSAMstrandField intronMotif --outSAMattributes All --outFilterMismatchNmax 10 --seedSearchStartLmax 30 --outSAMtype BAM SortedByCoordinate (Dobin et al., 2013). Then, subread (v1.5.2) was used to generate read counts data with the following parameters featureCounts -p -t exon -g gene_id (Liao et al., 2013). Significantly differentially expressed genes between treatments were analysed and selected using edgeR (v3.22.3) with false discovery rate (FDR) < 0.05 (Robinson et al., 2010). Removal of unwanted variance (RUVs) package in R was applied to two different batches of transcriptome datasets to eliminate batch variance (Risso et al., 2014). APE was used to cluster the treatments (Paradis et al., 2004) followed by RUVs. Gene Ontology (GO) overrepresentation analyses were performed using clusterProfiler with the parameter ont = "BP"(Biological Process), pAdjustMethod = "BH", pvalueCutoff = 0.01, and qvalueCutoff = 0.05 (Yu et al., 2012). Signalling Pathway Impact Analysis (SPIA) was carried out to identify the commonly perturbed pathways within the treatments using the SPIA R package (Tarca et al., 2008). Significantly perturbed pathways were visualized using Pathview package in R (Luo and Brouwer, 2013).

### Statistical analysis

Statistical analyses were carried out using GraphPad Prism 8.0 (GraphPad Software Inc., CA, USA). Student's t‐test or ANOVA (one‐way or two‐way) was used when there were two or three groups to compare respectively. Post hoc "Bonferroni's multiple comparisons test" was performed when ANOVA results were significant. Statistically significant results were represented as P<0.05 (*) or P<0.01 (**) P<0.001 (***), or P<0.0001 (****); ns (not significant). All data were shown as mean ± standard deviation (SD).

## Supporting information

## Acknowledgements

This work was supported by the special international corporation project of traditional Chinese medicine (GZYYGJ2017035) and The University of Adelaide, Zendong Australia China Centre for Molecular Chinese Medicine. The authors would like to thank Dr. Denis Scanlon and Associate Prof. Stephen Bell for the assistance with HPLC usage and Jue Zeng for her valuable discussions.

## Author contributions

T.N.A, S.N, Z.Q and D.L.A designed the study, analysed the data and wrote the manuscript. T.N.A and S.N conducted the experiments. Y.H-L, J.C, H.S, T.P, and H.D assisted the experiments and analysed the data. A.J.Y and D.K assisted in writing the manuscript.

## Conflict of interest

The authors declare no conflicts of interest.

## Declaration of transparency and scientific rigour

This declaration acknowledges that this paper adheres to the principles for transparent reporting and scientific rigour of preclinical research recommended by funding agencies, publishers and other organisations engaged with supporting research.

## Supplementary Figure 1

HPLC profiles of 25 mixtures including CKI, MJ, MN, 9 (N-1), 4 (N-3) and 9 (N-2). 50 <u>μ</u>l of the samples at 1 mg/ml concentration was injected through the semi-preparative column to achieve the profiles.

## Supplementary Figure 2

XTT Cell viability assays of subtractive fractions in MDA-MB-231 cells at 24- and 48-hour time points treated with 1 mg/ml or 2 mg/ml of CKI and 2 mg/ml equivalent concentrations of all other treating agents. (A) Suppression of cell viability from the following treatments: CKI, WRCKI-B (whole reconstituted CKI in buffer/vehicle control), and WRCKI-H (whole reconstituted CKI in milliQ H_2_O) (B) assessment of the interaction effects of single MJ compounds by the addition to the MN subtractive fraction. Single major compounds were dissolved in either MilliQ H_2_O or DMSO (Dimethyl sulfoxide). Effect of subtractive fractions (C) N-OmtOspcSpc, (D) N-MacAdeTri, (E) N-MtNmeSpr, (F) N-MtNme, (G) N-OmtSpc, (H) N-OspcSpc, (I) N-MacTri, (J) N-AdeTri, (K) N-MacAde, (L) N-MtSpr, (M) NmeSpr. Statistically significant results shown as P<0.05 (*) or P<0.01 (**) P<0.001 (***), or P<0.0001 (****); ns (not significant). All data were shown as mean ± standard deviation (SD).

## Supplementary Figure 3

XTT Cell viability assays of subtractive fractions N-OmtOspc and N-MacOmtOspc in MDA-MB-231 cells at 24- and 48-hour time points treated 4 different concentrations ranging from 0.25mg/mL to 2mg/mL of CKI and equivalent concentrations of two other agents. Statistically significant results shown as P<0.05 (*) or P<0.01 (**) P<0.001 (***), or P<0.0001 (****); ns (not significant). All data were shown as mean ± standard deviation (SD).

## Supplementary Figure 4

Representative plots of Annexin V and PI staining in (A) MDA-MB-231, (B) HEK-293, and (C) HFF.

## Supplementary Figure 5

Multiple dimensional scaling (MDS) plot for samples based on expression profiles of all genes before the removal of unwanted variance (RUVs in R package).

## Supplementary Figure 6

Multiple dimensional scaling (MDS) plot for samples based on expression profiles of all genes after the application of RUVs.

## Supplementary Figure 7

Box plot for samples based on expression profiles of all genes before the application of RUVs.

## Supplementary Figure 8

Box plot for samples based on expression profiles of all genes after the application of RUVs.

## Supplementary Figure 9

Venn diagrams showing the number of overlaping DE genes between treatments at 24-hours and 48-hours.

## Supplementary Figure 10

Identification of significantly perturbed pathways using SPIA (pG< 0.05) analysis. Eighty-six significantly perturbed pathways from twenty-two comparisons were found.

## Supplementary Figure 11

DE genes from the following comparisons (CKI vs UT, CKI vs N-Mac, CKI vs N-Nme, CKI vs N-Omt and CKI vs N-Tri) shown in the "Cytokine-Cytokine Receptor Interaction pathway at 48-hours. Significant up- and down-regulated DE genes were coloured red and green respectively. Each coloured box was separated into five parts according to treatments in this order: CKI vs UT, CKI vs N-Mac, CKI vs N-Nme, CKI vs N-Omt and CKI vs N-Tri. White or grey colours represented gene(s) that were not significantly differentially expressed by the treatments.

## Supplementary Figure 12

DE genes from the following comparisons (CKI vs UT, CKI vs N-OmtOspc, CKI vs N-MacOmtOspc, CKI vs OmtOspc and CKI vs MacOmtOspc) shown in the Cytokine-Cytokine Receptor Interaction pathway at 48-hours. Significant up- and down-regulated DE genes were coloured red and green respectively. Each coloured box was separated into five parts according to this order: CKI vs UT, CKI vs N-OmtOspc, CKI vs N-MacOmtOspc, CKI vs OmtOspc and CKI vs MacOmtOspc. White or grey colours represented gene(s) that were not significantly differentially expressed by the treatments.

## Supplementary Figure 13

DE genes from the following comparisons (CKI vs UT, CKI vs N-OmtOspc, CKI vs N-MacOmtOspc, CKI vs OmtOspc and CKI vs MacOmtOspc) shown in the Cytokine-Cytokine Receptor Interaction pathway at 24-hours. Significant up- and down-regulated DE genes were coloured red and green respectively. Each coloured box was separated into five parts according to this order: CKI vs UT, CKI vs N-OmtOspc, CKI vs N-MacOmtOspc, CKI vs OmtOspc and CKI vs MacOmtOspc. White or grey colours represented gene(s) that were not significantly differentially expressed by the treatments.

## Supplementary Figure 14

DE genes from the following comparisons (CKI vs UT, CKI vs N-Mac, CKI vs N-Nme, CKI vs N-Omt and CKI vs N-Tri) shown in the Cell Cycle pathway at 48-hours. Significant up- and down-regulated DE genes were coloured with red and green respectively. Each coloured box was separated into five parts according to this order: CKI vs UT, CKI vs N-Mac, CKI vs N-Nme, CKI vs N-Omt and CKI vs N-Tri. White or grey colours represented gene(s) that were not significantly differentially expressed by the treatments.

## Supplementary Figure 15

DE genes from the following comparisons (CKI vs UT, CKI vs N-OmtOspc, CKI vs N-MacOmtOspc, CKI vs OmtOspc and CKI vs MacOmtOspc) shown in the Cell Cycle pathway at 48-hour treatments. Significant up- and down-regulated DE genes were coloured red and green respectively. Each coloured box was separated into five parts according to this order: CKI vs UT, CKI vs N-OmtOspc, CKI vs N-MacOmtOspc, CKI vs OmtOspc and CKI vs MacOmtOspc. White or grey colours represented gene(s) that were not significantly differentially expressed by the treatments.

## Supplementary Figure 16

DE genes from the following comparisons (CKI vs UT, CKI vs N-OmtOspc, CKI vs N-MacOmtOspc, CKI vs OmtOspc and CKI vs MacOmtOspc) shown in the Cell Cycle pathway at 24-hours. Significant up- and down-regulated DE genes were coloured red and green respectively. Each coloured box was separated into five parts according to this order: CKI vs UT, CKI vs N-OmtOspc, CKI vs N-MacOmtOspc, CKI vs OmtOspc and CKI vs MacOmtOspc. White or grey colours represented gene(s) that were not significantly differentially expressed by the treatments.

## Supplementary Figure 17

Differential gene expression profiles of all treatments for TGF-ß signalling pathway: the left panel shows comparison of subtractive fraction treated cells against CKI treatment and the right panel shows comparison of single compound subtractive fraction treated cells against the treatments for two and three compound subtractive fractions.

## Supplementary Figure 18

DE genes from the following comparisons (CKI vs UT, CKI vs N-Mac, CKI vs N-Nme, CKI vs N-Omt and CKI vs N-Tri) shown in the TGF-ß signalling pathway at 48-hours. Significant up- and down-regulated DE genes were coloured red and green respectively. Each coloured box was separated into five parts according to this order: CKI vs UT, CKI vs N-Mac, CKI vs N-Nme, CKI vs N-Omt and CKI vs N-Tri. White or grey colours represented gene(s) that were not significantly differentially expressed by the treatments.

## Supplementary Figure 19

DE genes from the following comparisons (CKI vs UT, CKI vs N-OmtOspc, CKI vs N-MacOmtOspc, CKI vs OmtOspc and CKI vs MacOmtOspc) shown in the TGF-ß signalling pathway at 48-hour treatments. Significant up- and down-regulated DE genes were coloured red and green respectively. Each coloured box was separated into five parts according to this order: CKI vs UT, CKI vs N-OmtOspc, CKI vs N-MacOmtOspc, CKI vs OmtOspc and CKI vs MacOmtOspc. White or grey colours represented gene(s) that were not significantly differentially expressed by the treatments.

## Supplementary Figure 20

DE genes from the following comparisons (CKI vs UT, CKI vs N-OmtOspc, CKI vs N-MacOmtOspc, CKI vs OmtOspc and CKI vs MacOmtOspc) shown in the TGF-ß signalling pathway at 24-hours. Significant up- and down-regulated DE genes were coloured red and green respectively. Each coloured box was separated into five parts according to this order: CKI vs UT, CKI vs N-OmtOspc, CKI vs N-MacOmtOspc, CKI vs OmtOspc and CKI vs MacOmtOspc. White or grey colours represented gene(s) that were not significantly differentially expressed by the treatments.

## Notes

#### Summary of Updates

Updated discussion.

