## Supplementary material for "Fractional Deletion Of Compound Kushen Injection, A Natural Compound Mixture, Indicates Cytokine Signaling Pathways Are Critical For Its Perturbation Of The Cell Cycle": Supplementary Figure 1.pdf

**Supplementary Figure 1**: HPLC profiles of 25 mixtures including CKI, MJ, MN, 9 (N-1), 4 (N-3) and 9 (N-2). 50uL of the samples at 1mg/mL concentration was injecteded through the semi-preparative column to achieve the profiles.
