## Supplementary material for "Fractional Deletion Of Compound Kushen Injection, A Natural Compound Mixture, Indicates Cytokine Signaling Pathways Are Critical For Its Perturbation Of The Cell Cycle": SupplFigure16_CellCycle_CKIvsUTN23OOMOO_24.pdf

Treatments (24hrs, from left to right) : CKI vs UT, CKI vs N-OmtOspc, N-MacOmtOspc, CKI vs OmtOspc, CKI vs MacOmtOspc

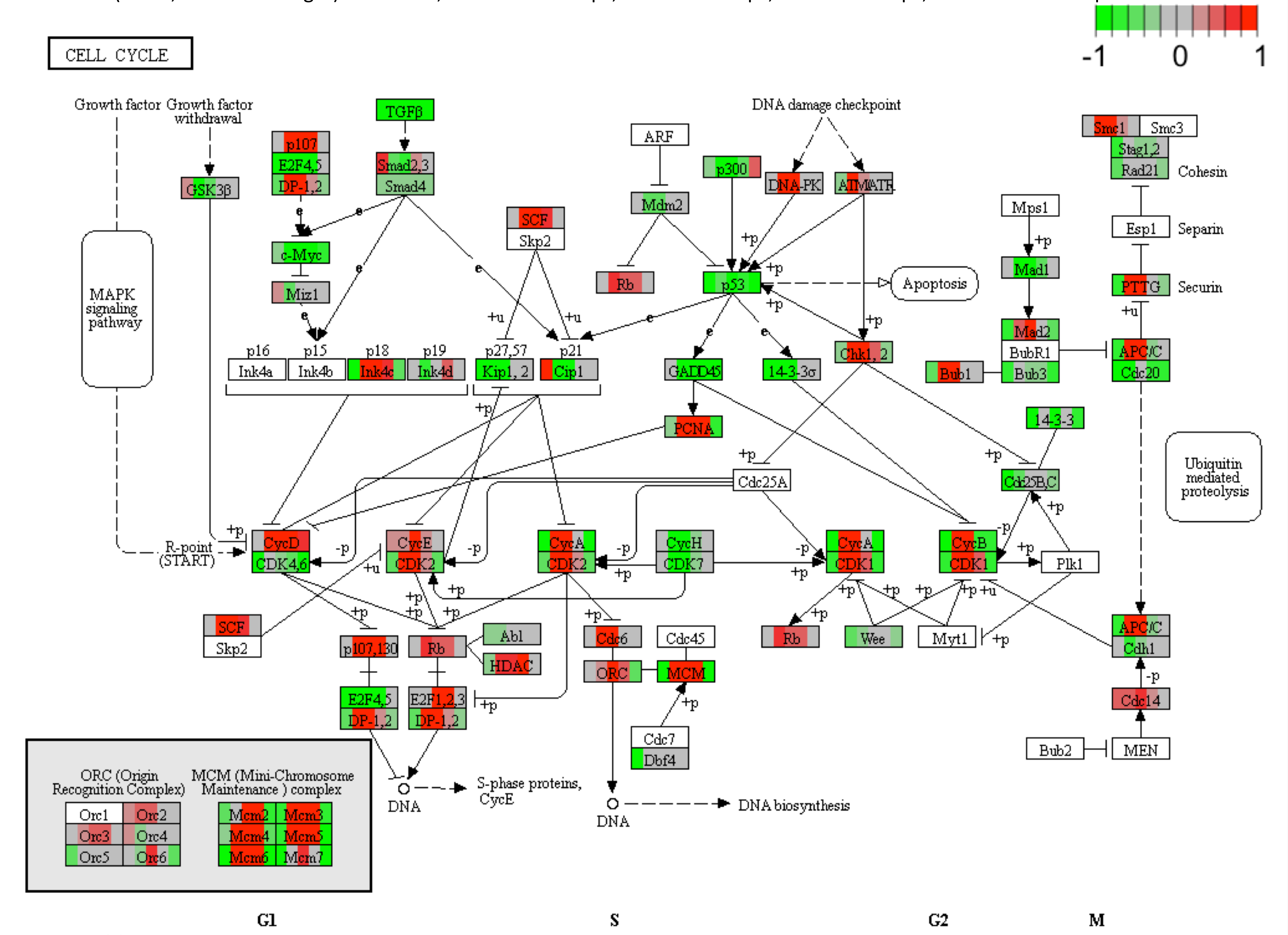


Data on KEGG graph
Rendered by Pathview
