## Supplementary material for "Fractional Deletion Of Compound Kushen Injection, A Natural Compound Mixture, Indicates Cytokine Signaling Pathways Are Critical For Its Perturbation Of The Cell Cycle": SupplFigure18_TGF_beta_CKIvsN-1_UTMacNmeOmtTri_.pdf

Treatments (48hrs, from left to right) : CKI vs UT, CKI vs N-Mac, CKI vs N-Nme, CKI vs N-Omt, CKI vs N-Tri

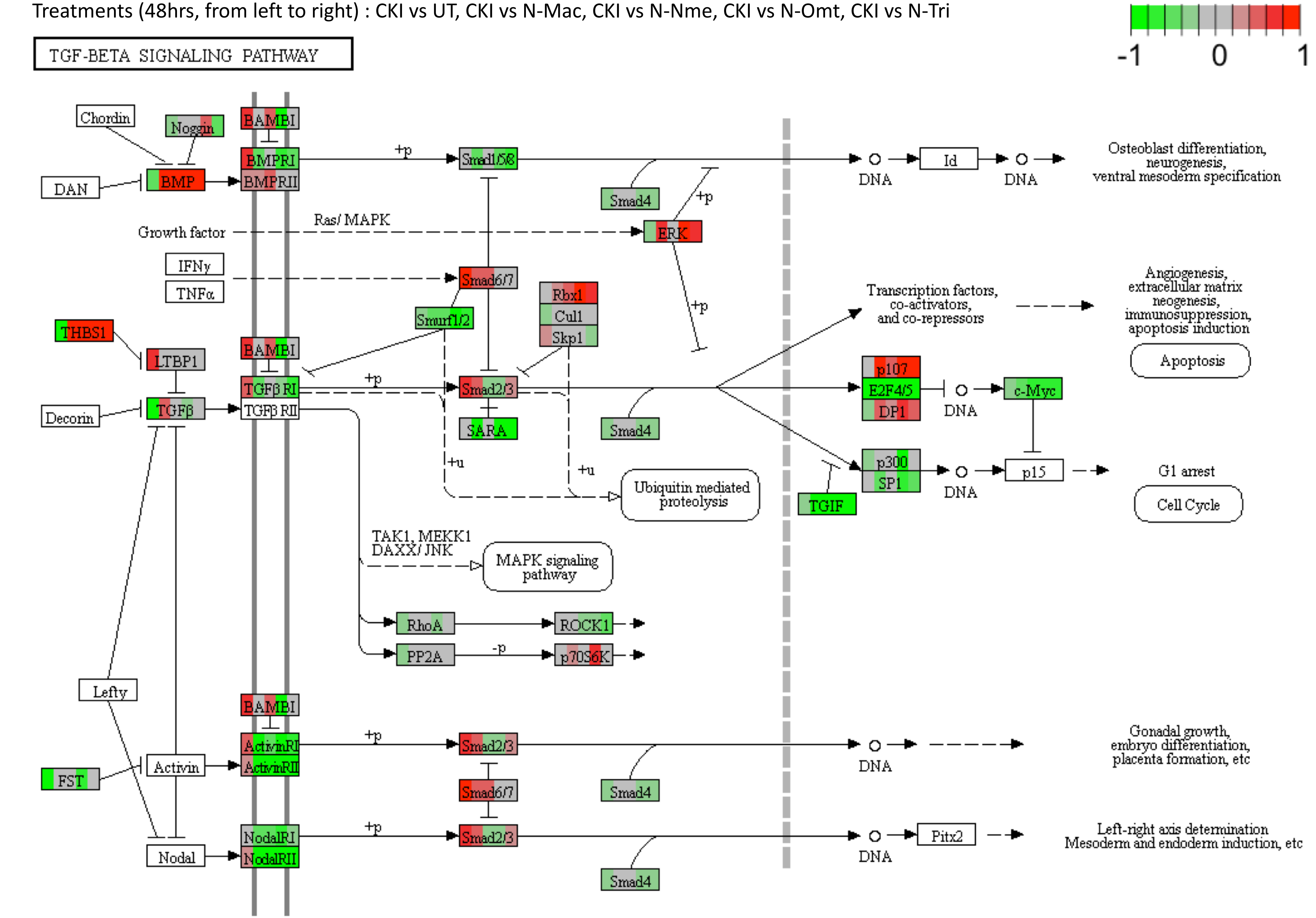


Data on KEGG graph
Rendered by Pathview
