## Supplementary material for "Fractional Deletion Of Compound Kushen Injection, A Natural Compound Mixture, Indicates Cytokine Signaling Pathways Are Critical For Its Perturbation Of The Cell Cycle": SupplFigure19_TGF_CKIvsUTN23OOMOO_48.pdf

Treatments (48hrs, from left to right) : CKI vs UT, CKI vs N-OmtOspc, N-MacOmtOspc, CKI vs OmtOspc, CKI vs MacOmtOspc

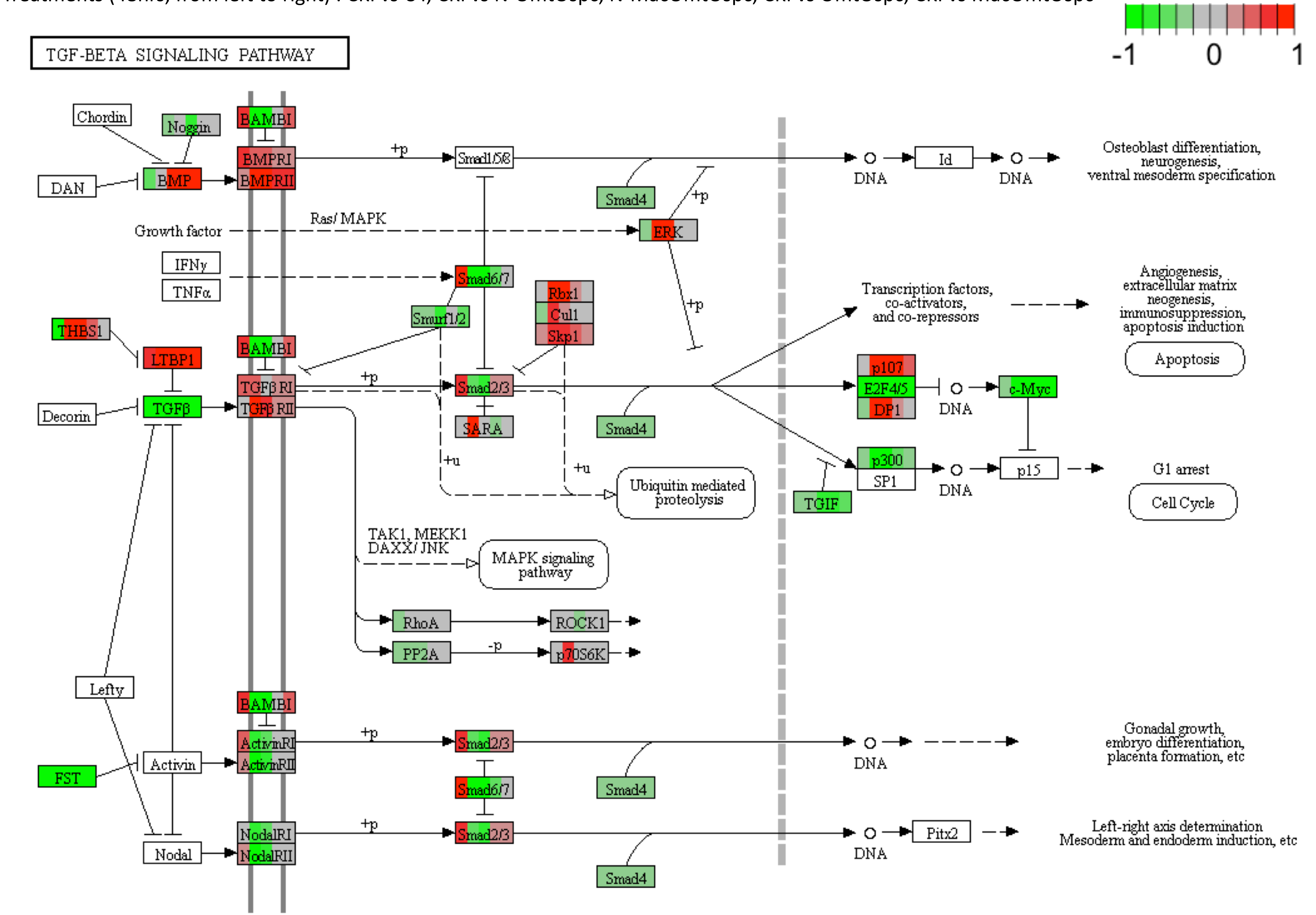


Data on KEGG graph
Rendered by Pathview
