## Supplementary material for "Fractional Deletion Of Compound Kushen Injection, A Natural Compound Mixture, Indicates Cytokine Signaling Pathways Are Critical For Its Perturbation Of The Cell Cycle": SupplFigure20_TGF_CKIvsUTN23OOMOO_24.pdf

Treatments (24hrs, from left to right) : CKI vs UT, CKI vs N-OmtOspc, N-MacOmtOspc, CKI vs OmtOspc, CKI vs MacOmtOspc

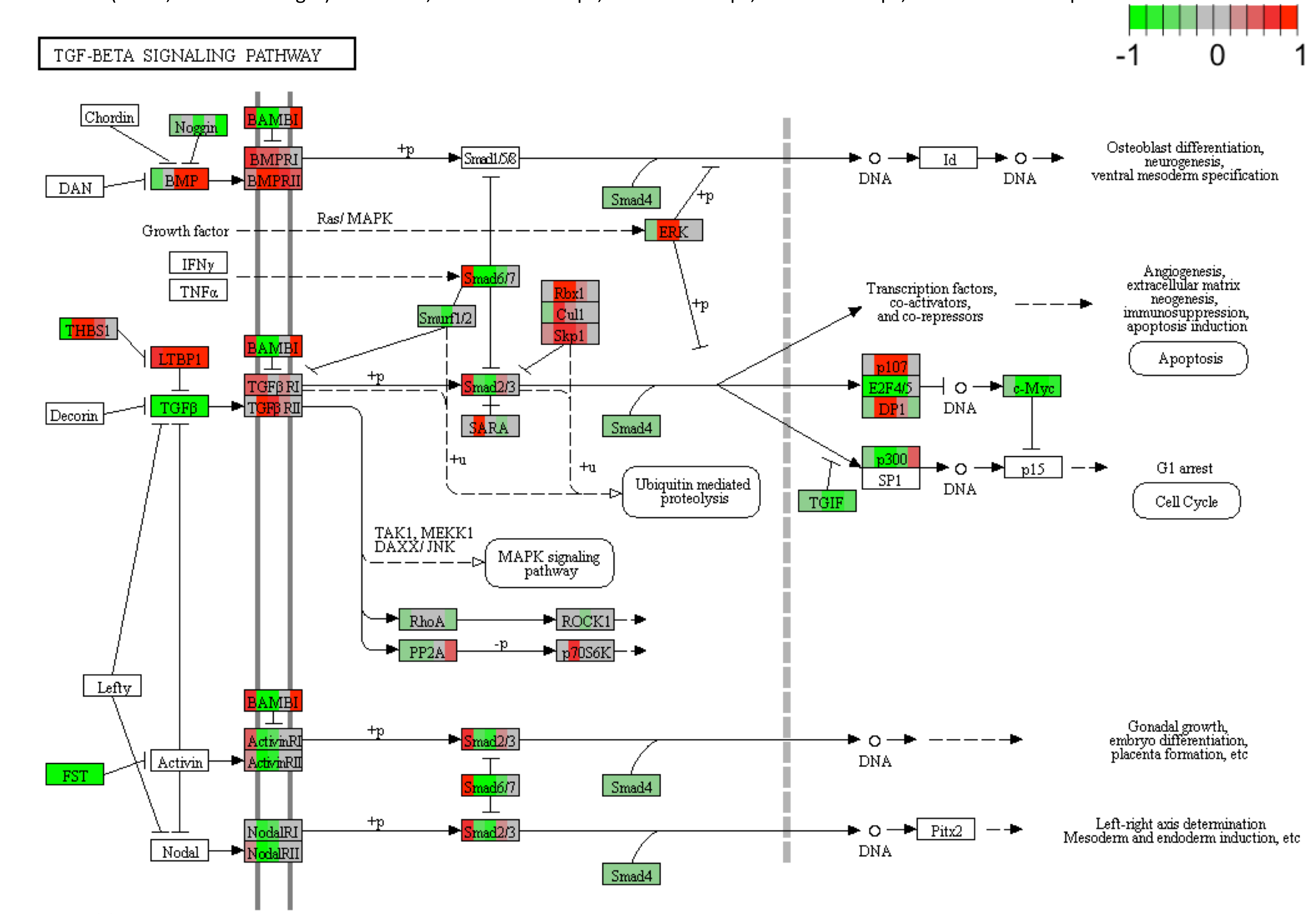


**Data on KEGG graph**
**Rendered by Pathview**
