## Supplementary figures and images for "Fractional Deletion Of Compound Kushen Injection, A Natural Compound Mixture, Indicates Cytokine Signaling Pathways Are Critical For Its Perturbation Of The Cell Cycle"

### SupplFigure2.pdf

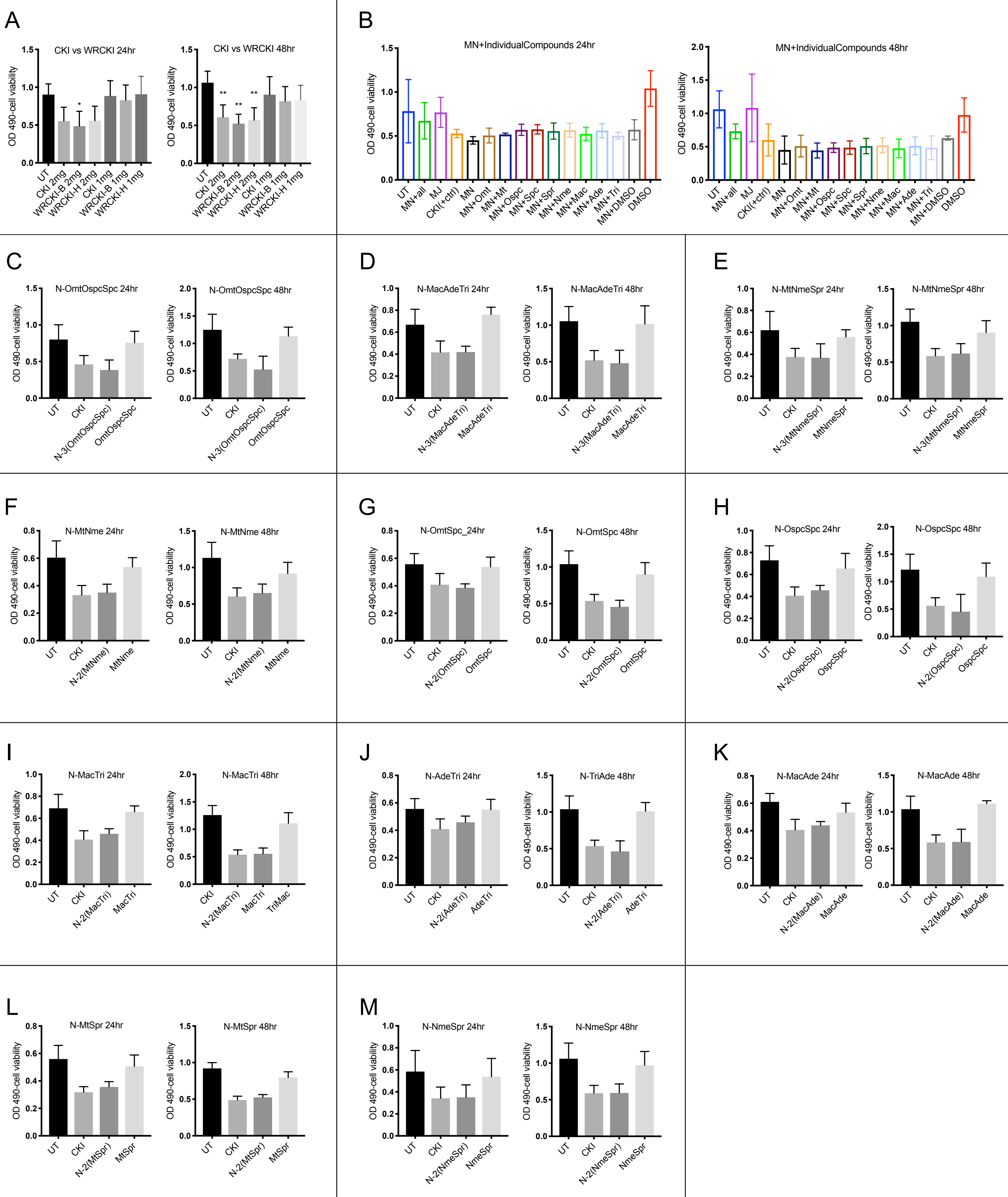

### SupplFigure3.pdf

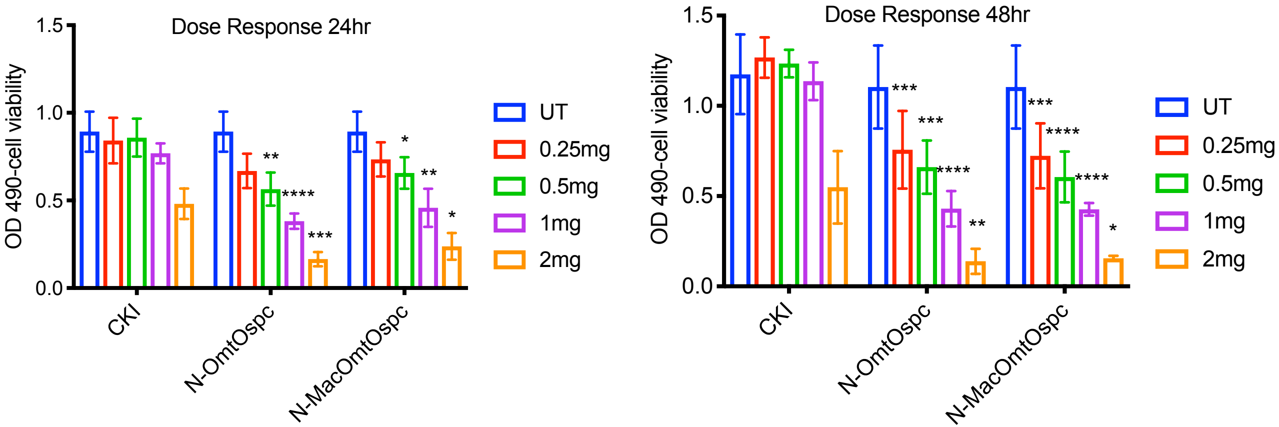

### SupplFigure4.pdf

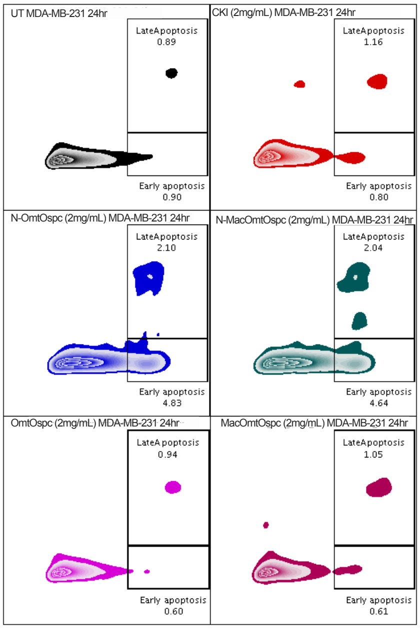

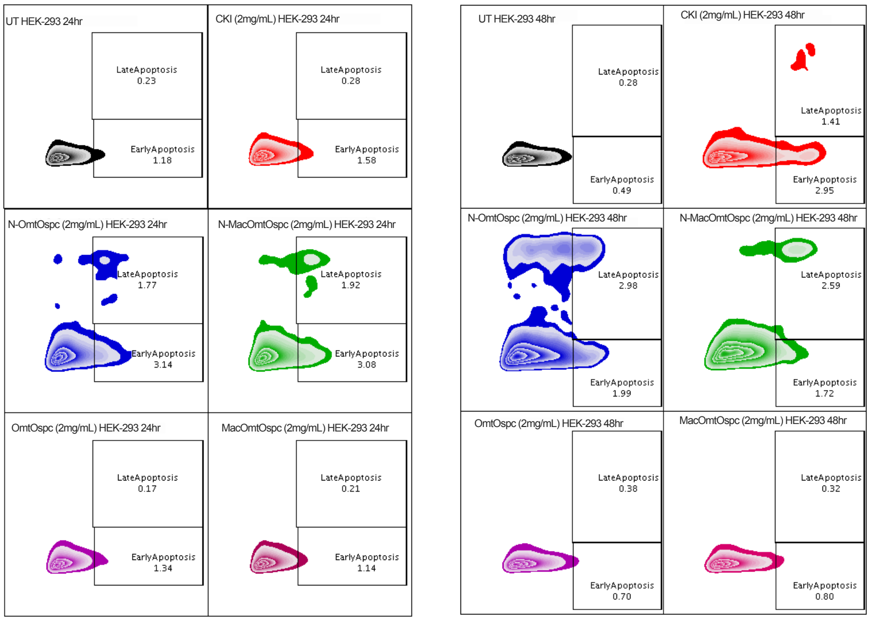

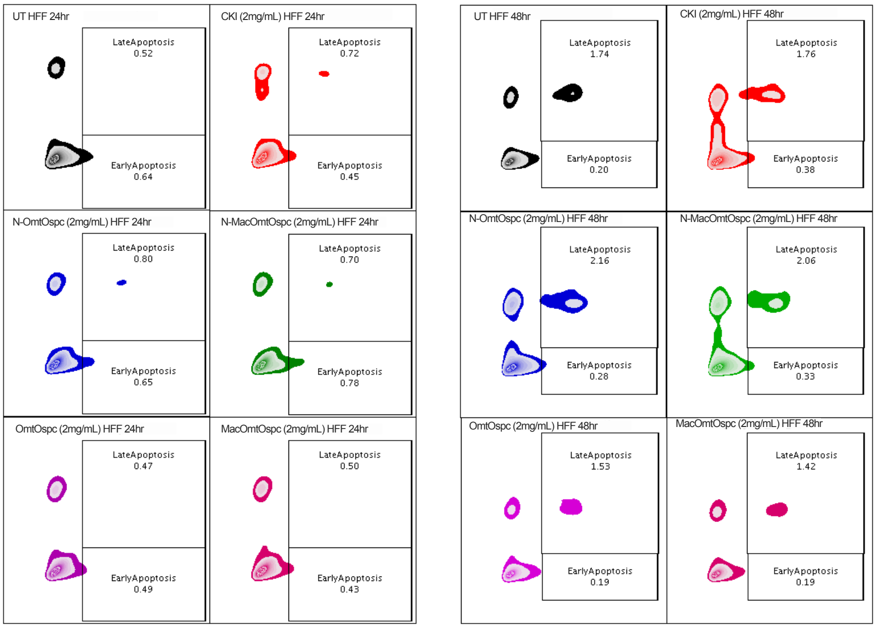

### SupplFigure5_MDS.pdf

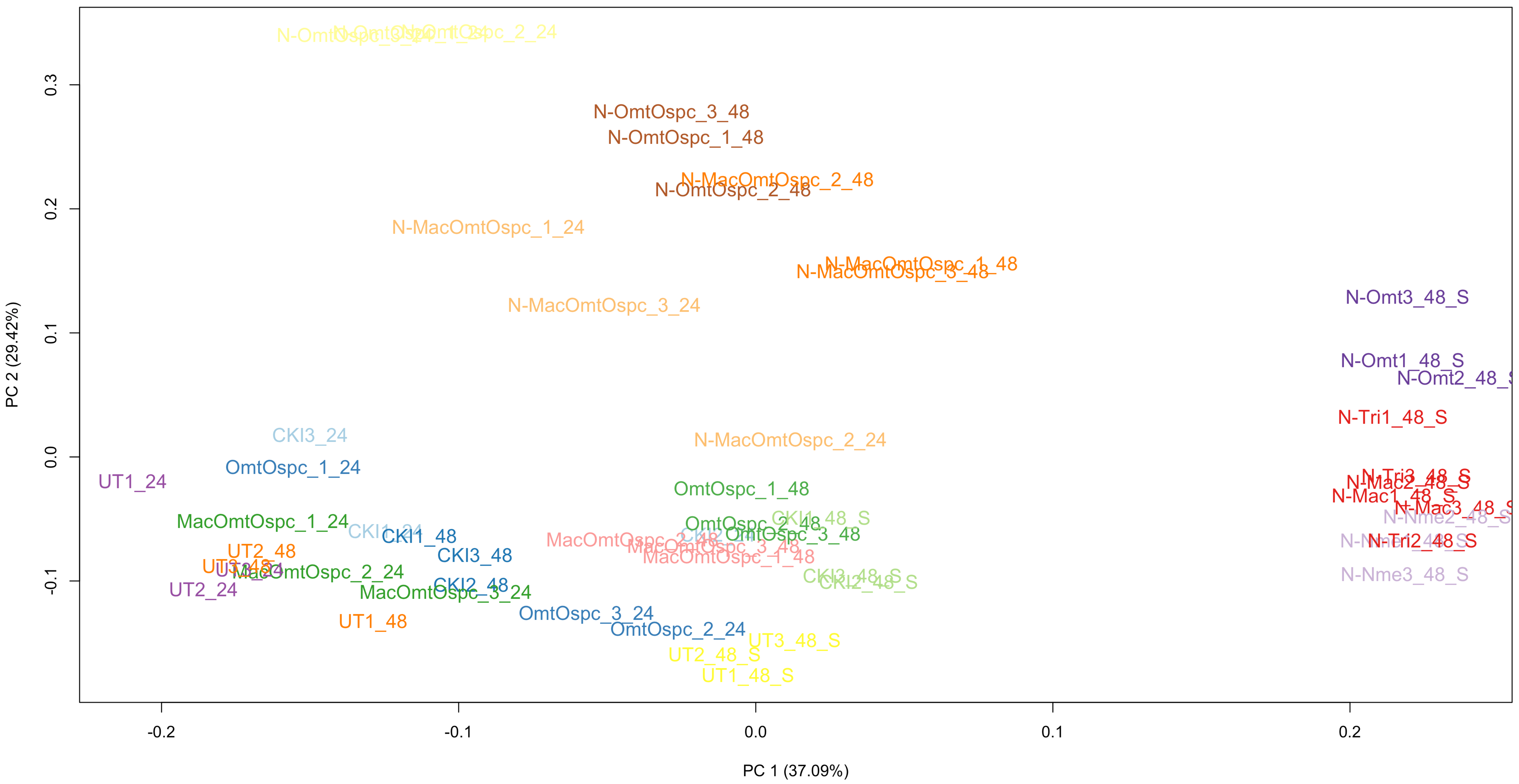

### SupplFigure6_MDS.pdf

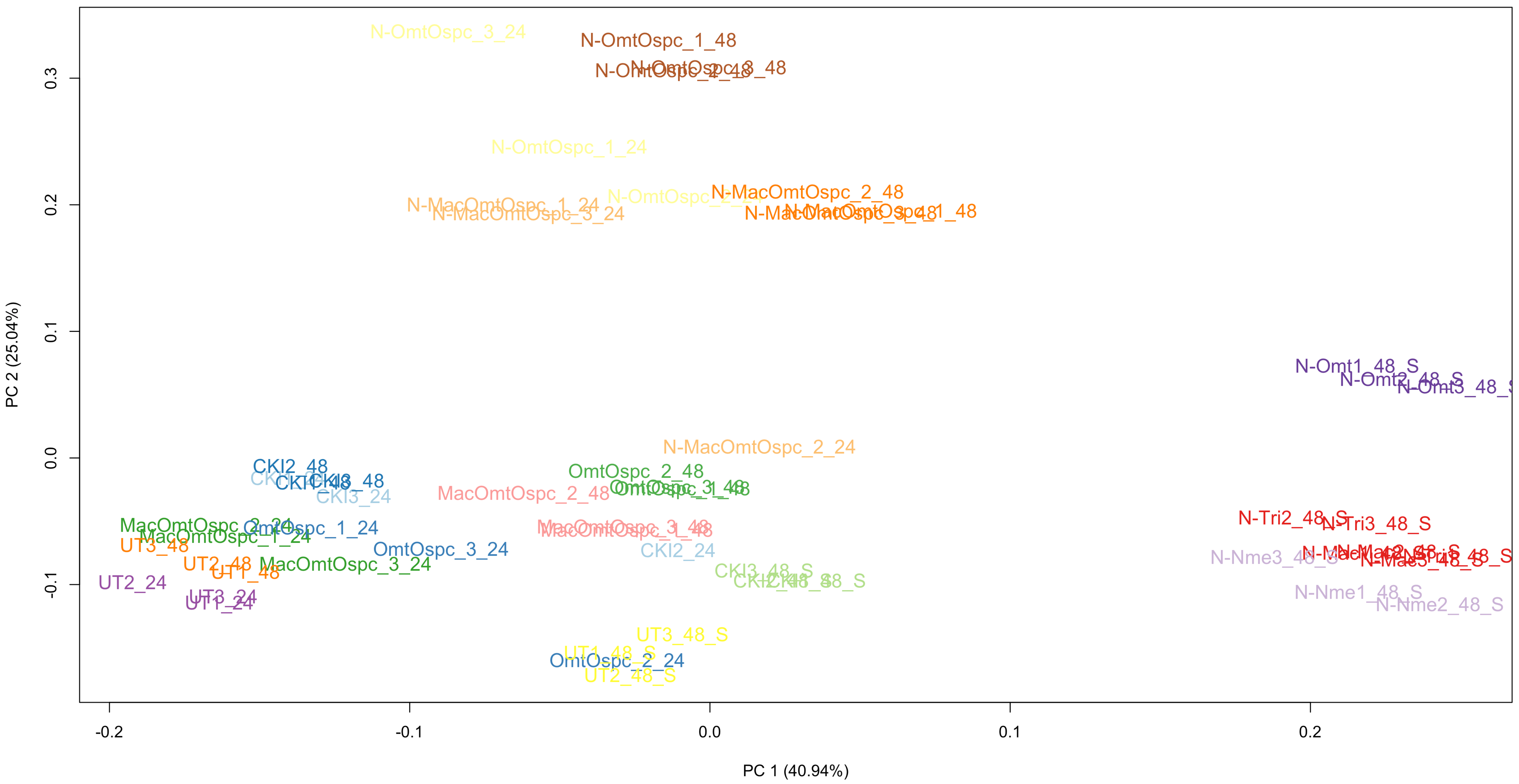

### SupplFigure7_RUV.pdf

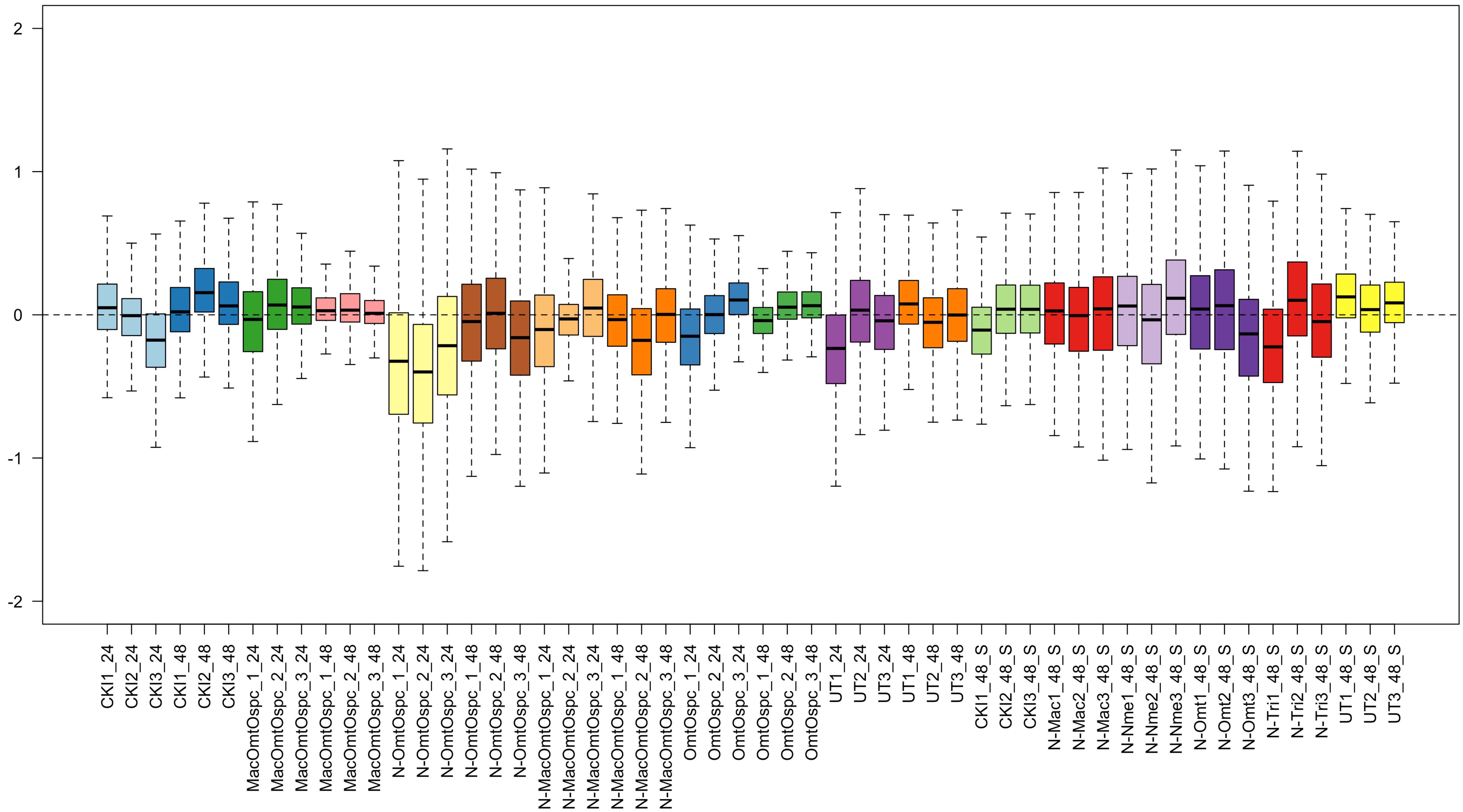

### SupplFigure8_RUV.pdf

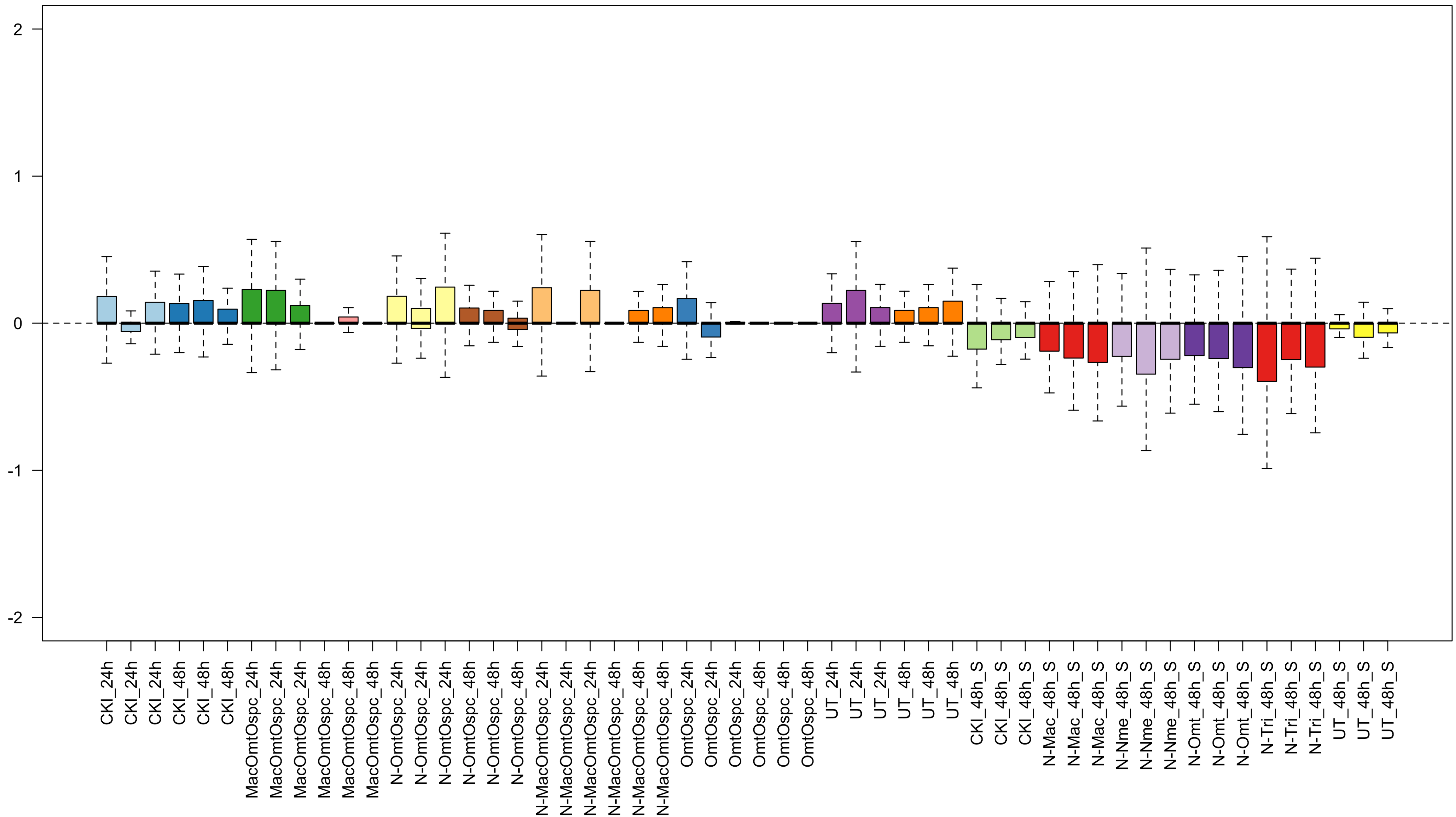

### SupplFigure9_venn_24_48.pdf

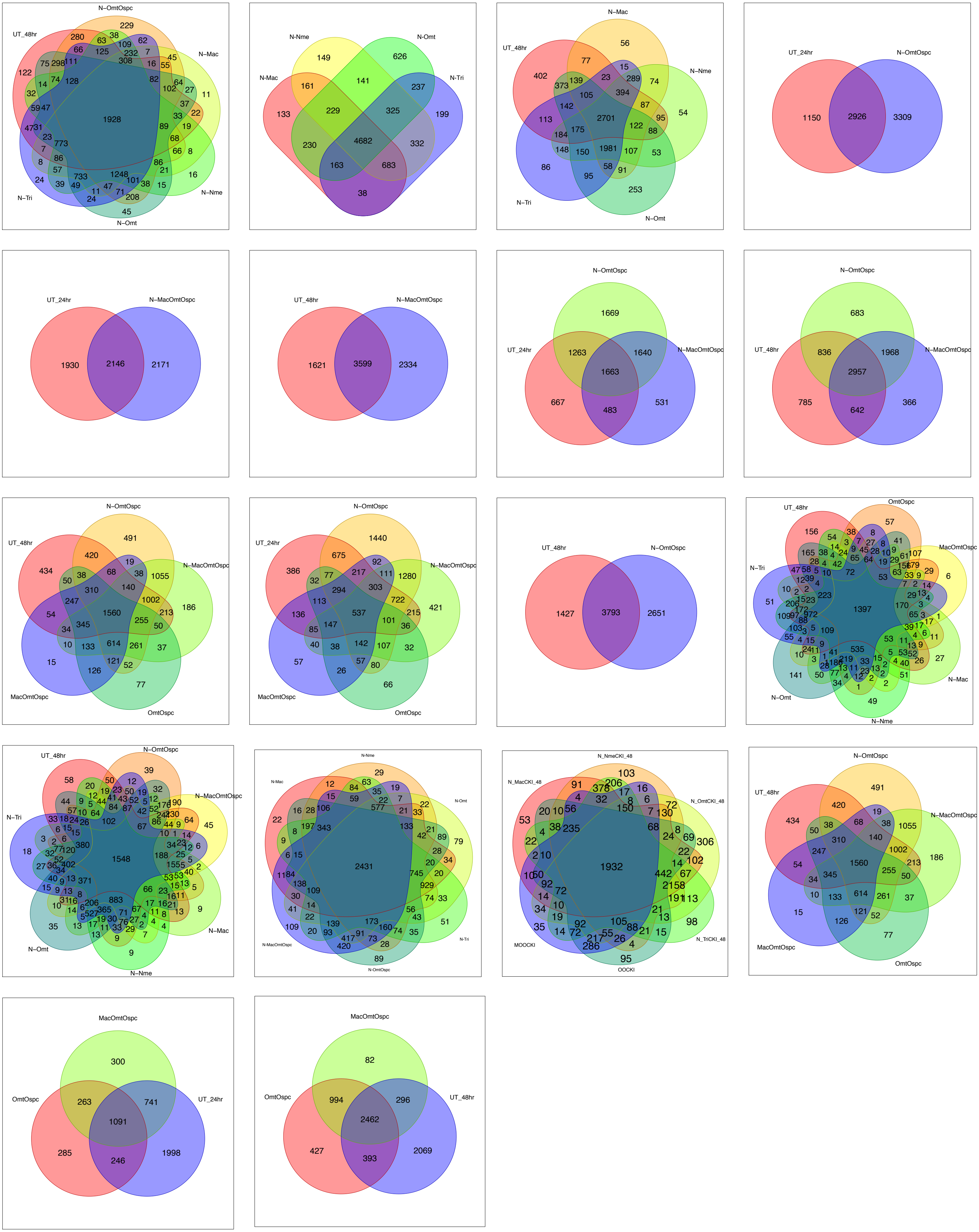

### SupplFigure10_SPIA.pdf

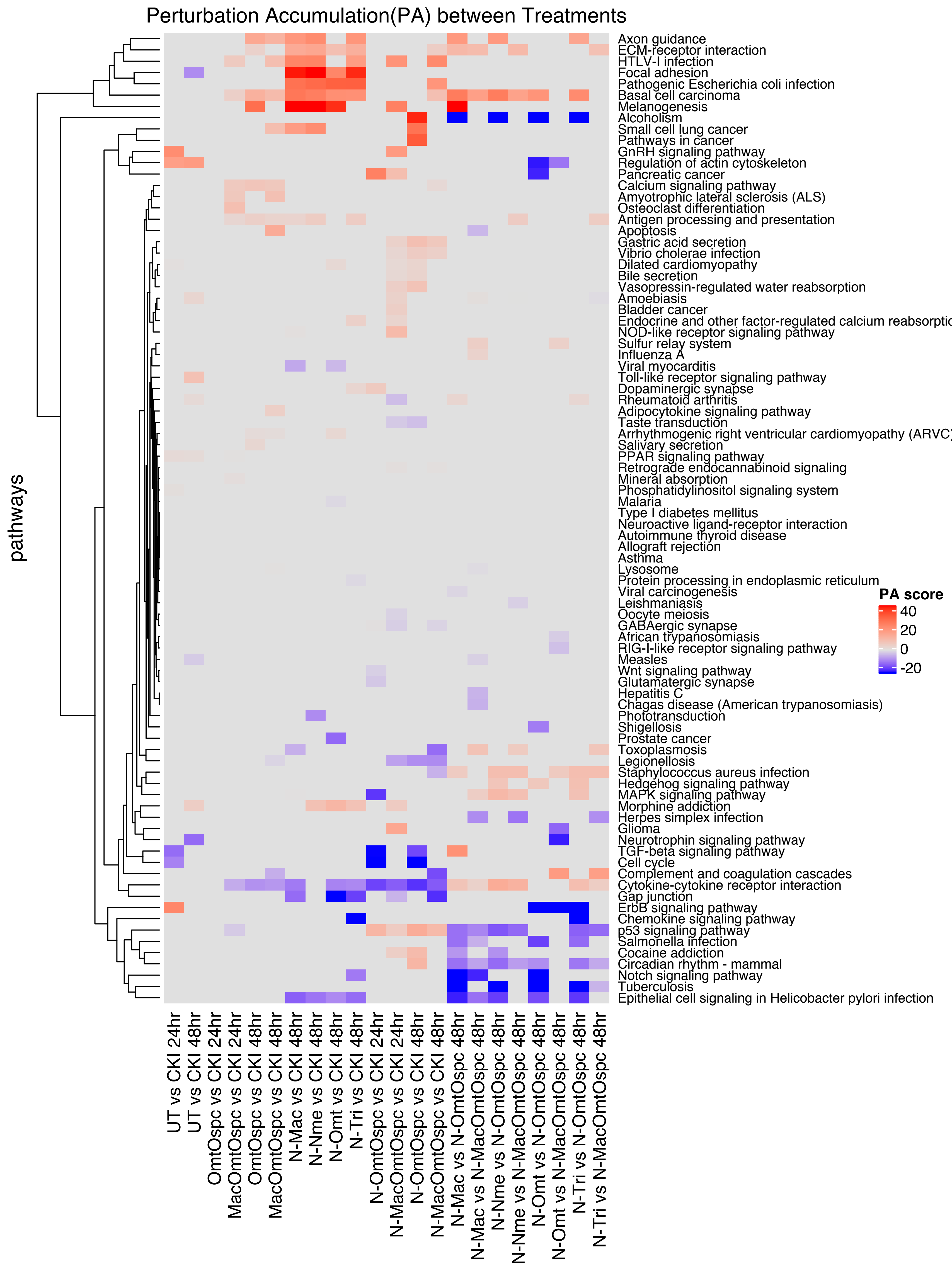

### SupplFigure11_Cytokine_CKIvsN-1_UTMacNmeOmtTri_.pdf

Treatments (48hrs, from left to right) : CKI vs UT, CKI vs N-Mac, CKI vs N-Nme, CKI vs N-Omt, CKI vs N-Tri

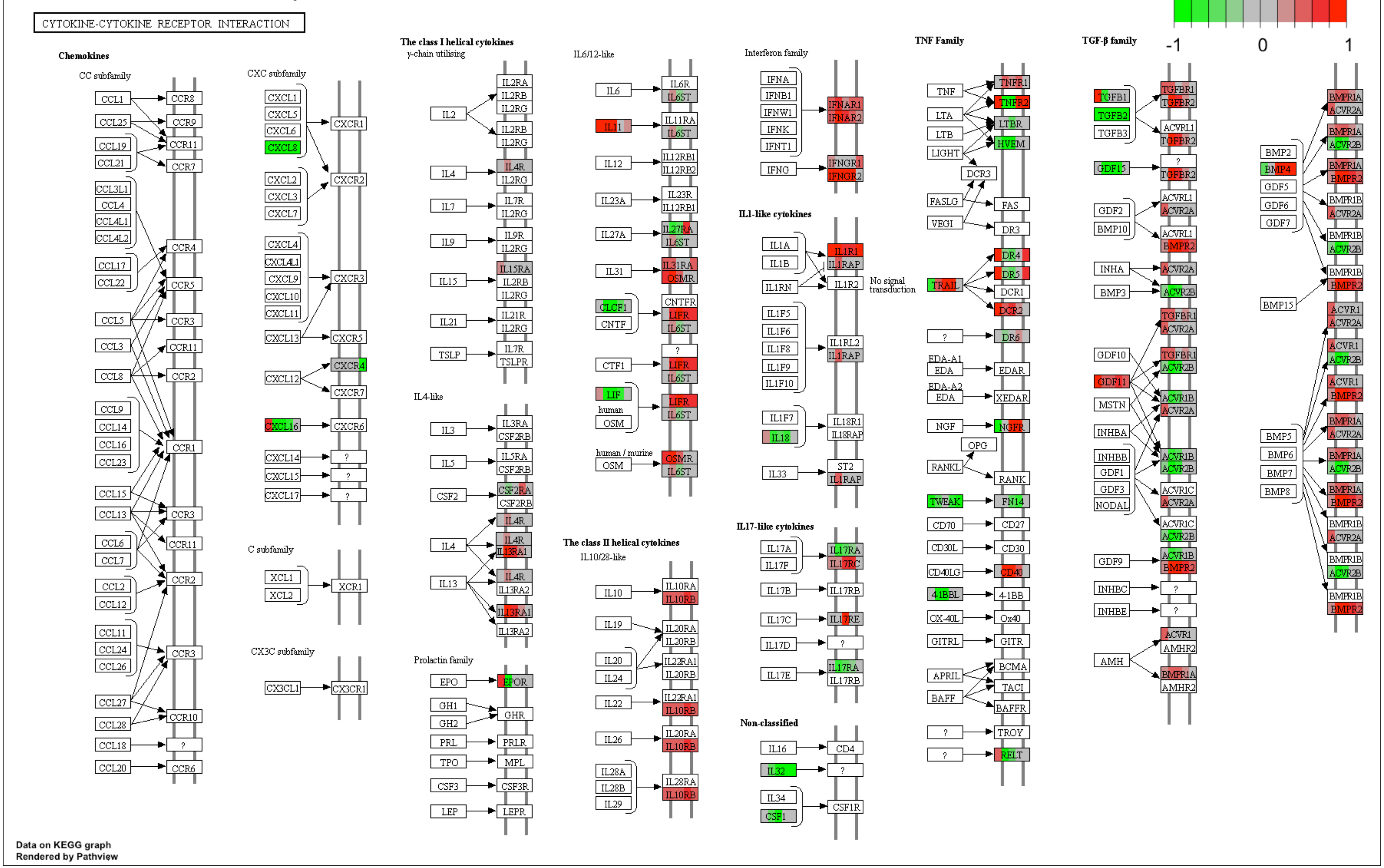

### SupplFigure14_CellCycle_CKIvsN-1_UTMacNmeOmtTri_.pdf

Treatments (48hrs, from left to right) : CKI vs UT, CKI vs N-Mac, CKI vs N-Nme, CKI vs N-Omt, CKI vs N-Tri

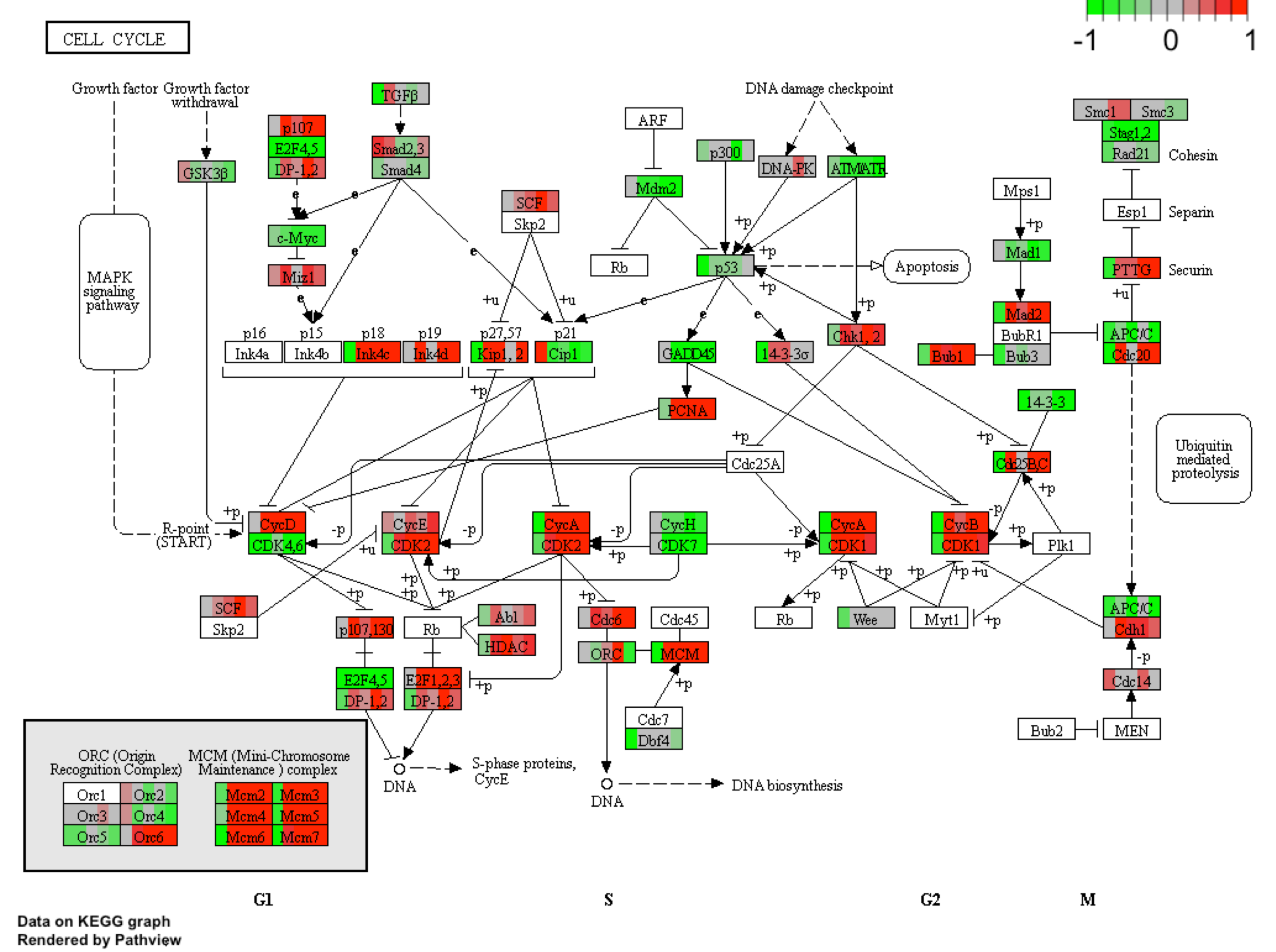

### SupplFigure17TGF_betaSignaling_2112018.pdf

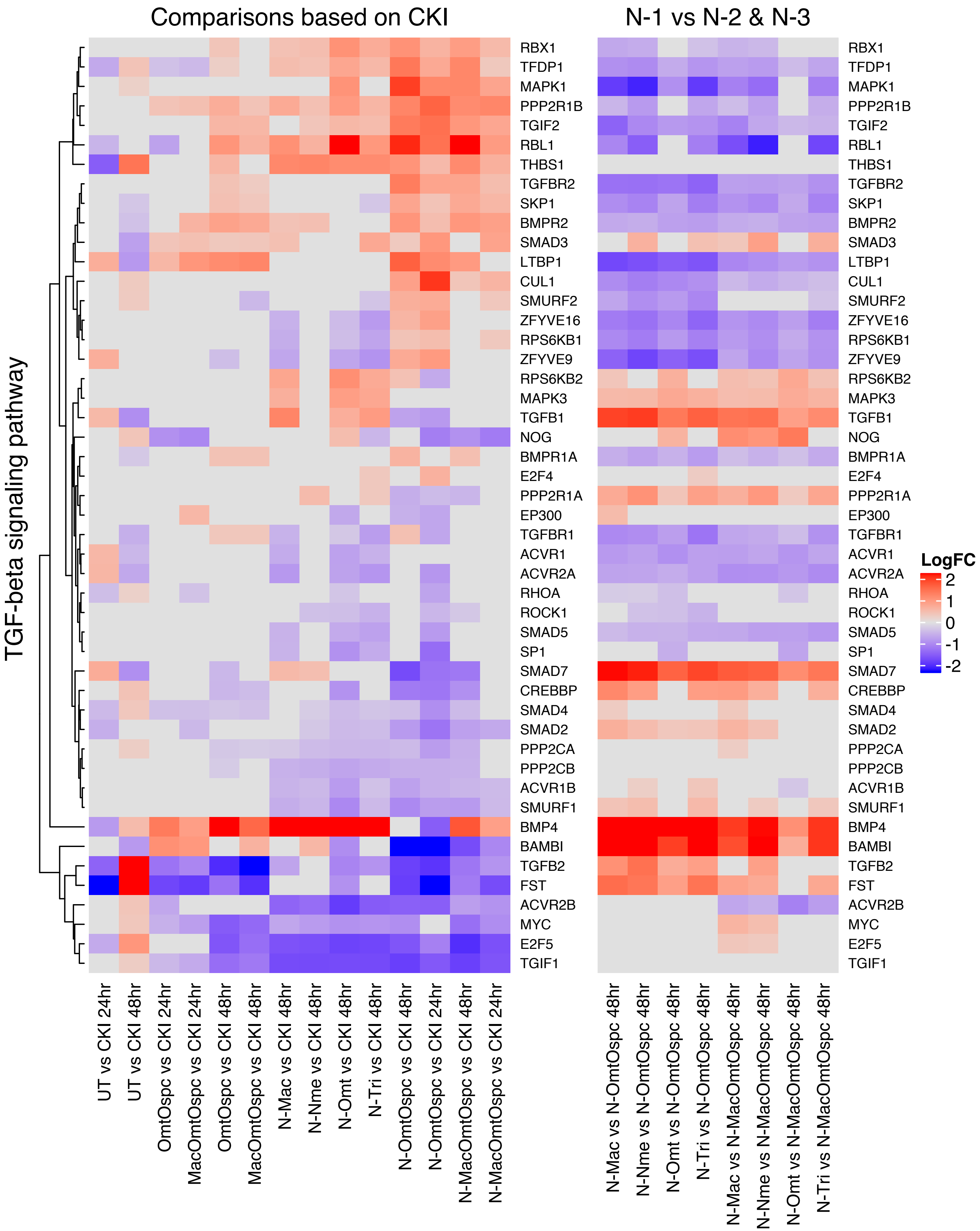
